# Bacterial Geosmin Biosynthesis is Compartmentalized Inside a Two-Component Protein Shell

**DOI:** 10.64898/2026.07.21.739788

**Authors:** Asif Fazal, Cassandra A. Dutcher, Michael P. Andreas, Tobias W. Giessen

## Abstract

Terpenoids are ubiquitous, structurally diverse natural products found across the three domains of life. One of the most commonly observed and well-studied terpenoids is the volatile, petrichoric compound geosmin, responsible for the earthy smell of soil to which humans are receptive down to a few parts per trillion. Bioinformatic analysis of bacterial geosmin synthase (GeoA)-encoding gene clusters reveals that many are colocalized with genes encoding Family 2B encapsulin shell proteins. Here, we focus on the encapsulin-encoding geosmin gene cluster of the model myxobacterium *Myxococcus xanthus* and show that the two shell proteins form a two-component Family 2B encapsulin with GeoA as the internalized cargo. Structural analysis highlights that the shell contains characteristic, external cyclic adenosine monophosphate (cAMP) binding-fold domains (CBDs), and that mixed shells can adopt a previously unobserved closed two-fold pore conformation, potentially important in cargo function modulation and regulation. We show that GeoA cargo loading is mediated by repeating, short cargo loading peptides (CLPs), and that GeoA encapsulation provides benefits for enzyme activity and stability. This work expands the short list of Family 2B encapsulins known to be involved in specialized metabolite biosynthesis, and provides insights into a putative mode of cargo protein regulation via pore state modulation.

## Introduction

Terpenoids, also called isoprenoids, represent the most expansive and structurally diverse family of natural products, with over 80,000 compounds identified across the three domains of life [1-3]. Despite their immense structural complexity, all terpenoids are derived from two five-carbon (*C*_5_) isoprene units: isopentenyl pyrophosphate (IPP) and its isomer dimethylallyl pyrophosphate (DMAPP) [4]. Through the action of polyprenyl transferases, terpene synthases and subsequent tailoring enzymes, such as cytochrome P450 monooxygenases, these simple building blocks are transformed into a vast array of linear and cyclic scaffolds [5, 6]. These structures serve as the foundation for essential biological molecules, including sterols, carotenoids, and various plant and bacterial secondary metabolites [7, 8]. Iterative additions of IPP to the DMAPP starter unit results in the formation of the various acyclic terpenoid precursors. Geranyl pyrophosphate (GPP, *C*_10_) is the starting point for the synthesis of monoterpenoid products while farnesyl pyrophosphate (FPP, *C*_15_) and geranylgeranyl pyrophosphate (GGPP, *C*_20_) lead to sesqui- and diterpenoids, respectively [9].

Historically, terpenoids have been isolated from plants and fungi, but the recent expansion in availability of genomic data has shown that bacteria encode many natural product biosynthetic gene clusters (BGCs) containing polyprenyl transferases and terpene synthases and, accordingly, have the potential to be prodigious producers of terpenoids [1, 10]. Two of the most prevalent terpenoids produced by bacteria are 2-methylisoborneol (2-MIB) and geosmin [11, 12]. Both geosmin and 2-MIB are produced predominantly by soil-dwelling actinobacteria and myxobacteria, but production has also been observed in cyanobacteria and fungal species [13, 14]. Although the precise biological functions of these compounds are not known, recent hypotheses have suggested roles in productive attraction or as a predator repellant [15, 16]; specifically, the volatile nature of these compounds, coupled with the timing of production in *Streptomyces* coinciding with the onset of sporulation, suggest a potential role in spore dispersal by insect attraction [12, 17]. Interestingly, both geosmin and 2-MIB are non-standard terpenoids, with the *C*_11_ 2-MIB formed from a methylated GPP precursor and the *C*_12_ geosmin formed from cyclization of the sesquiterpene precursor FPP followed by the loss of a *C*_3_ acetone unit.

Geosmin biosynthesis has been studied extensively since the isolation of the compound in 1965 and the determination of its chemical structure a few years later [18, 19]. Biosynthetic analyses have centered around the enzyme GeoA, which was observed to catalyze the conversion of the fifteen carbon FPP precursor to geosmin through a two-step process [20, 21]. GeoA is composed of two distinct terpene cyclase domains connected by a long flexible linker [22]. The initial on-pathway reaction to geosmin resembles a classical terpene cyclization, with C-C bond formation and concomitant loss of pyrophosphate leading to the production of the intermediate compound germacradienol in a Mg^2+^ dependent manner (Figure 1A, Figure S1) [23, 24]. Geosmin formation is then accomplished by an unprecedented retro-Prins fragmentation, catalyzed by the GeoA C-terminal domain [25]. Isotopic labelling studies and mutational analyses support this route to geosmin, as well as the production of the minor side products germacrene D and octalin (Figure S1) [26, 27].

**Figure 1:**
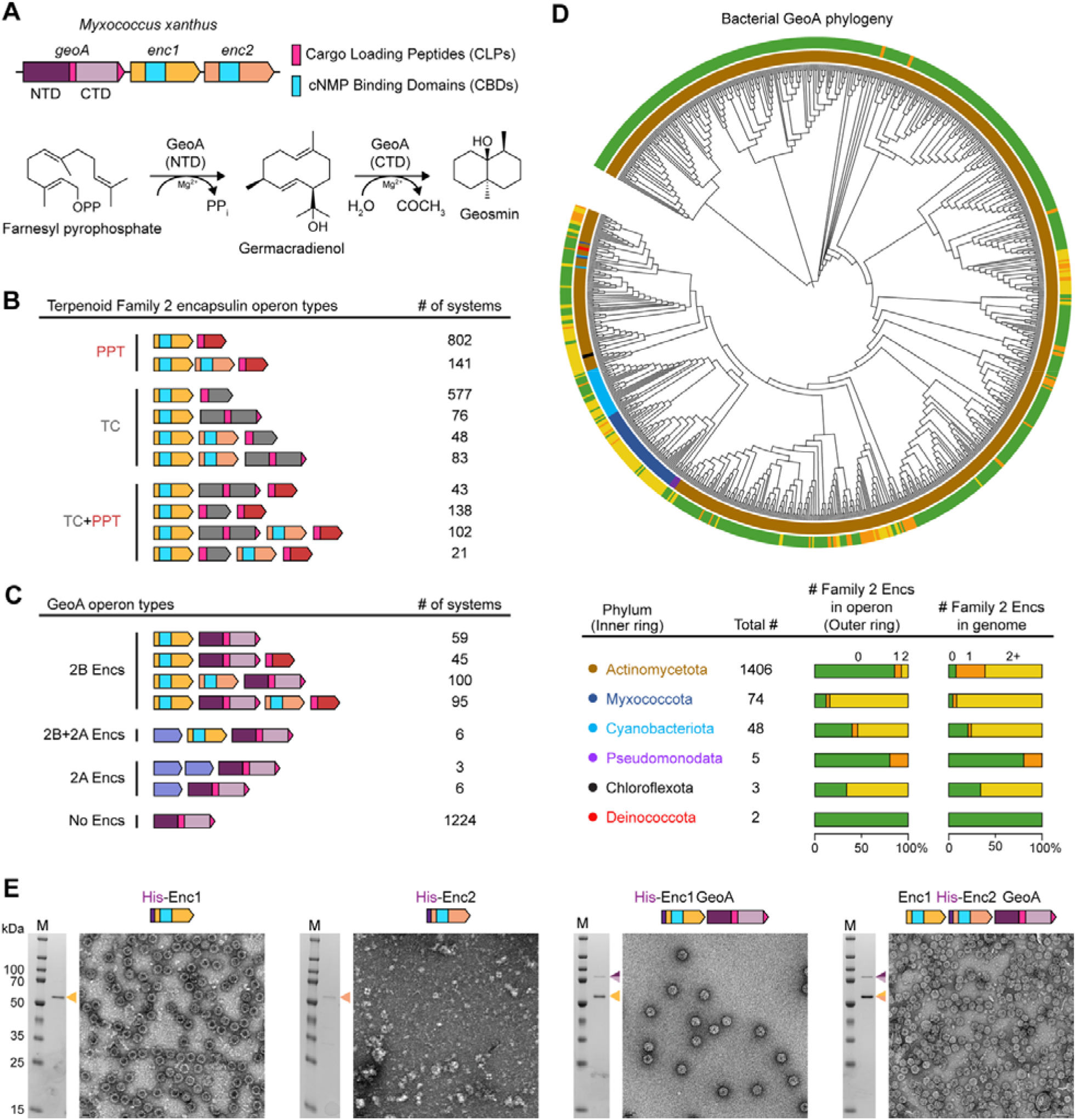
Gene clusters encoding encapsulin systems and terpene biosynthesis, including geosmin biosynthesis are widespread in bacteria. (A) The geosmin biosynthetic gene cluster from *Myxococcus xanthus*, encoding two family 2B encapsulins and GeoA, the biosynthetic enzyme required for geosmin formation. Cargo Loading Peptides (CLPs) and cNMP Binding Domains (CBDs) are indicated, as well as the N-terminal domain (NTD) and C-terminal domain (CTD) of GeoA. Biosynthesis of geosmin, by GeoA from farnesyl pyrophosphate (FPP) is also shown. (B) An overview of the encapsulin-associated terpene biosynthetic systems. Family 2B encapsulins were found to be associated with polyprenyl transferase (PPT) or terpene cyclase (TC) enzymes, or with a combination of both biosynthetic proteins. (C) Overview of all GeoA homologs observed in a separate bioinformatic analysis. The number of each type of system observed is noted beside each operon type. (D) Phylogenetic analysis of all observed GeoA homologs. The inner ring outside of the tree corresponds to the GeoA-associated phylum, with colors indicated in the table. The outer ring corresponds to the number of encapsulin genes observed in the GeoA-associated operon, with colors indicated in the table. The table also shows the total number of GeoA homologs found in each lineage as well as the number of encapsulins found in each operon or associated genome, respectively. (E) SDS-PAGE and negative stain transmission electron microscopy (TEM) analysis of proteins purified from each of the noted *E. coli* based expression strains.

Encapsulin nanocompartments (encapsulins) are a class of prokaryotic protein compartments, widely observed across bacterial and archaeal phyla [28, 29]. Generally, encapsulins are self-assembling, cargo-loaded protein shells, where the nature of the protein cargo dictates the specific functionality of the encapsulin-cargo system; roles in metal storage, stress resistance, natural product biosynthesis, and elemental metabolism have been observed to date [30-33]. All encapsulin shell proteins share the HK97 phage-like fold and form icosahedral protein shells that range in size from ca. 20 to 50 nm in diameter, with triangulation numbers of T = 1 (60 subunits), T = 3 (180 subunits) or T = 4 (240 subunits) [34, 35]. The eponymous feature of encapsulins is their ability to sequester and encapsulate specific cargo proteins, a process mediated by targeting or cargo loading sequences at the N- or C-termini of native cargos [36]. Encapsulins are characterized into four families based on sequence similarity of the encapsulin genes and the level of conservation observed in their genome neighborhoods [28]. Family 1 encapsulins are the most well-studied, whilst Family 2 is subdivided into two classes, 2A and 2B, based on the absence or presence of putative cyclic nucleotide-binding domains (CBDs) within the encapsulin shell protein [37-40]. A Family 2B encapsulin has recently been shown to encapsulate the 2-MIB synthase 2-MIBS [41], whilst another Family 2B encapsulin shell was observed to be formed from two distinct encapsulin protomers, highlighting the diversity and variability of Family 2B systems [42].

Here, using the *Myxococcus xanthus* system, we show that GeoA is internalized as cargo within a two-component Family 2B encapsulin shell. The encapsulated GeoA shows an unhindered kinetic profile, whilst encapsulation also provides a proteolysis-prevention phenotype compared to the free enzyme. Cryo-electron microscopy (cryo-EM) studies of the encapsulin shell show that cargo-loading is mediated through extended, flexible cargo loading motifs, and that the presence of the second encapsulin protomer in shells leads to altered pore conformations with the potential to modulate internal enzyme activity. Overall, this work further expands the roles played by Family 2B encapsulins in terpenoid biosynthesis and gives initial indications into how two-component encapsulin shells may control catalysis by modulating lumenal flux of substrate and product molecules.

## Results and Discussion

### Geosmin Biosynthetic Gene Clusters Encode Family 2B Encapsulin Genes

Recent studies have highlighted that multiple terpene and terpenoid BGCs also contain genes encoding for Family 2B encapsulin shells [30, 33]. Further, our recent study on the biosynthesis of 2-MIB showed that in multiple bacterial phyla, the primary biosynthetic enzyme 2-MIB synthase, is encapsulated inside a Family 2B shell [41]. Our further analysis of Family 2B encapsulin genome neighborhoods revealed that many contain genes encoding for putative terpenoid biosynthetic genes, with 2425 systems observed in total encoding either a polyprenyl transferase (PPT) or a terpene cyclase (TC), or both types of enzymes (Figure 1B). Intriguingly, of these putatively Family 2B encapsulated terpenoid-forming systems, 299 (12.3%) were found to contain a TC resembling GeoA, the two-domain enzyme catalyzing geosmin formation (Figure 1C) [43, 44]. Further, 65% of these GeoA-containing operons encode two distinct encapsulin genes, indicating the potential for mixed shell formation. It is also noteworthy that of the 1224 GeoA-containing systems that do not exhibit a colocalized encapsulin-encoding gene, the vast majority (88.3%) are present in organisms that encode at least one Family 2B encapsulin elsewhere in the genome (Figure 1D).

Phylogenetic analysis of bacterial GeoA homologs shows that the majority (91%) are of actinobacterial origin, with most of the remainder encoded within Myxococcota and Cyanobacteriota (Figure 1D). Actinobacteria preferentially encode *geoA* genes without a neighboring encapsulin, whereas the majority of Myxococcota and Cyanobacteriota coencode *geoA* with putative shell forming genes, potentially indicating that geosmin biosynthesis can be encapsulated in these organisms. In this study, we focus on the GeoA- and encapsulin-encoding BGC from *Myxococcus xanthus* (Mx) as a pertinent case study for the occurrence and role of encapsulation within geosmin biosynthesis.

### Variable Combinations of Family 2A and Family 2B Encapsulin Genes Colocalize with GeoA

The *geoA* gene within *M. xanthus* is encoded alongside two putative Family 2B encapsulin-encoding genes, henceforth called *MxEnc1* and *MxEnc2*, producing proteins MxEnc1 and MxEnc2 (Figure 1A). The Mx genome, as mentioned, is one of many that encodes GeoA alongside Family 2B encapsulins. Further inspection of bacterial GeoA homologs, however, also revealed the presence of numerous systems containing either one or two Family 2A encapsulins, as well as *geoA* colocalization with both Family 2A and Family 2B shell genes (Figure 1C). While one-component Family 2A encapsulin systems were observed in Actinobacteria and *Pseudomonas* species, certain actinobacterial genomes were found to contain different two-component systems (either 2x Family 2A or 1x Family 2A and 1x Family 2B) (Figure S2).

The vast majority of Family 2A encapsulins are associated with cysteine desulfurase cargos and not terpenoid-related cargo proteins [31, 32, 45]. Indeed, no Family 2A systems encoding TCs have been experimentally characterized, and evidence of their existence has only been observed at the genomic level in a handful of instances [29]. The observation of putative GeoA-type TC encapsulation within a Family 2A system is intriguing as most TCs are Family 2B shell encapsulated. Family 2B shells exhibit open pore structures and the potential for further post-translational and post-encapsulation enzyme regulation, via pore size modulation enabled by the presence of CBDs, as hypothesized previously [35, 46]. Family 2A systems would have to follow a different molecular logic for enzyme regulation. We further observed six actinobacterial genomes encoding systems containing a *geoA* homologue and both a Family 2A and Family 2B shell gene, with other putative biosynthetic enzymes encoded within the BGC as well (Figure S2). These hybrid 2A/2B systems have not been observed previously, and raise the possibility of altered encapsulation logic of either separate Family 2A and 2B shells formed by the same BGC, or a two-component hybrid encapsulin formed from both 2A and 2B protomers. An example system showing this type of BGC organization is found in *Nonomuraea* sp. C10, which contains both a PPT and GeoA-like TC for putative terpenoid biosynthesis (Figure S2). Sequence and AlphaFold-based analyses show that all encapsulin protomers adopted the classical HK97-like fold seen in all encapsulin systems [47]. Analysis of putative cargo proteins revealed that all GeoAs contain extended C-termini of varying lengths, as well as a disordered stretch of residues between the two TC domains. Both disordered regions could act as cargo loading peptides (CLPs) mediating cargo encapsulation [48, 49].

### The *M. xanthus* Geosmin Gene Cluster Encodes a Two-Component Family 2B Encapsulin With GeoA as its Cargo Protein

*In silico* analysis of the geosmin gene cluster of Mx showed the presence of two neighboring Family 2B encapsulin-encoding genes directly downstream of *geoA* (Figure 1A, S1). The Mx GeoA enzyme (MxGeoA) has 55% identity (70% similarity) to the characterized *Streptomyces coelicolor* GeoA, indicating that MxGeoA likely catalyzes the same two step conversion of FPP to germacradienol, through to the final geosmin product (Figure S3) [21]. The effect of the presence of the encapsulin shell on geosmin biosynthesis has not been characterized before in any system.

We first sought to investigate whether MxEnc1 and MxEnc2 are able to form single- or two-component protein shells, and if so, if GeoA could be cargo-loaded. A previous study on two-component encapsulin systems showed that one protomer was able to form shells by itself, whereas the second component produced from the system did not form larger shell-resembling complexes when produced in isolation [42]. When both protomers were produced in tandem, the result was the production of mixed shells with stochastically incorporated protomers, where no specific protomer-protomer interactions were seemingly disallowed.

In order to observe their propensity to form encapsulin shells, we heterologously expressed and purified an N-terminally His-tagged variant of each protomer in *Escherichia coli*. Negative stain transmission electron microscopy (TEM) showed that His-MxEnc1 formed clear regular shells with a diameter of ca. 27 nm, consistent with other T = 1 encapsulin shells (Figure 1E) [31, 32, 41]. Protrusions on the exterior of the shell could also be observed by TEM, likely representing externally displayed CBDs [35]. Purification and TEM analysis of His-MxEnc2, however, yielded no observable shell formation and no other regular macromolecular assemblies (Figure 1E). As discussed, this is in agreement with observations from other two-component Family 2B shells, where only one protomer in isolation can form discrete encapsulin shells [42].

We then moved to investigating whether cargo-loaded shells could be formed from the Mx system. To do this, we coexpressed Mx *geoA* with either one or both encapsulins. MxGeoA clearly copurified with His-MxEnc1 and analysis by TEM again showed the presence of T = 1 encapsulin shells, this time with more interior electron dense puncta, likely representing the encapsulated MxGeoA proteins. The same results were also obtained when MxGeoA was coexpressed with both His-MxEnc1 and His-MxEnc2, with TEM showing encapsulin shell formation (Figure 1E). SDS-PAGE could not resolve the two encapsulin bands but the presence of both protomers was confirmed by mass spectrometry analysis, following in-gel tryptic digest (Figure S1). To further clarify any ambiguity surrounding shell composition, and the presence of both protomers within shells when MxEnc1 and MxEnc2 were copurified, we generated a strain expressing MxEnc1 and MxGeoA with His-MxEnc2. As His-MxEnc2 cannot form shells alone, any affinity-purified shells must represent mixed MxEnc1/His-MxEnc2 shells. Purification of encapsulin shells from this strain again showed coelution of encapsulin and GeoA cargo bands, with TEM imaging showing the same T = 1 shell architecture (Figure 1E). This experiment strongly suggests that GeoA-loaded mixed MxEnc1/His-MxEnc2 shells were formed.

### Cryo-EM Analysis of GeoA-Loaded Encapsulins Highlights Overall Shell Architecture and the Presence of Open and Closed Pore States in Mixed Shells

In order to further investigate the molecular structure of the Mx Family 2B encapsulin system, we carried out single-particle cryo-EM analysis on both the one-component, cargo-loaded shell (His-MxEnc1_GeoA), and a two-component cargo-loaded sample (MxEnc1_His-MxEnc2_GeoA) (Figure 2). Icosahedral refinements yielded very similar consensus cryo-EM maps for both samples with global resolutions of 2.31 Å and 2.27 Å, respectively (Figure S4, S5). Neither asymmetric reconstructions nor masked classifications allowed us to differentiate MxEnc1 from MxEnc2 protomers in mixed shells, likely due to their high sequence similarity (79%; sequence identity: 63%) and stochastic arrangement. The I-symmetry averaged mixed density closely resembled the MxEnc1-only map. The shells each consist of 60 protomers arranged with T = 1 icosahedral symmetry. This is consistent with our TEM analysis and with previous reports of Family 2B encapsulin structures, which displayed the same triangulation pattern (Figure 2A) [41, 42]. Lower resolution CBDs were visible at the exterior of the shell, arranged symmetrically around all two-fold axes of symmetry. In addition, extra non-protomer density could be observed located at interior three-fold axes of symmetry, representing GeoA CLPs. The pores at each of the two-fold axes were large and extended in nature, with dimensions measuring ∼20 x 60 Å (Figure S6). These extended pores sit underneath, but are not occluded by, the external CBDs and their size is characteristic of Family 2B encapsulin shells (Figure 2B). The five-fold and three-fold pores are much smaller by comparison, with sizes of ∼6 Å and ca. ∼5 Å, respectively.

**Figure 2:**
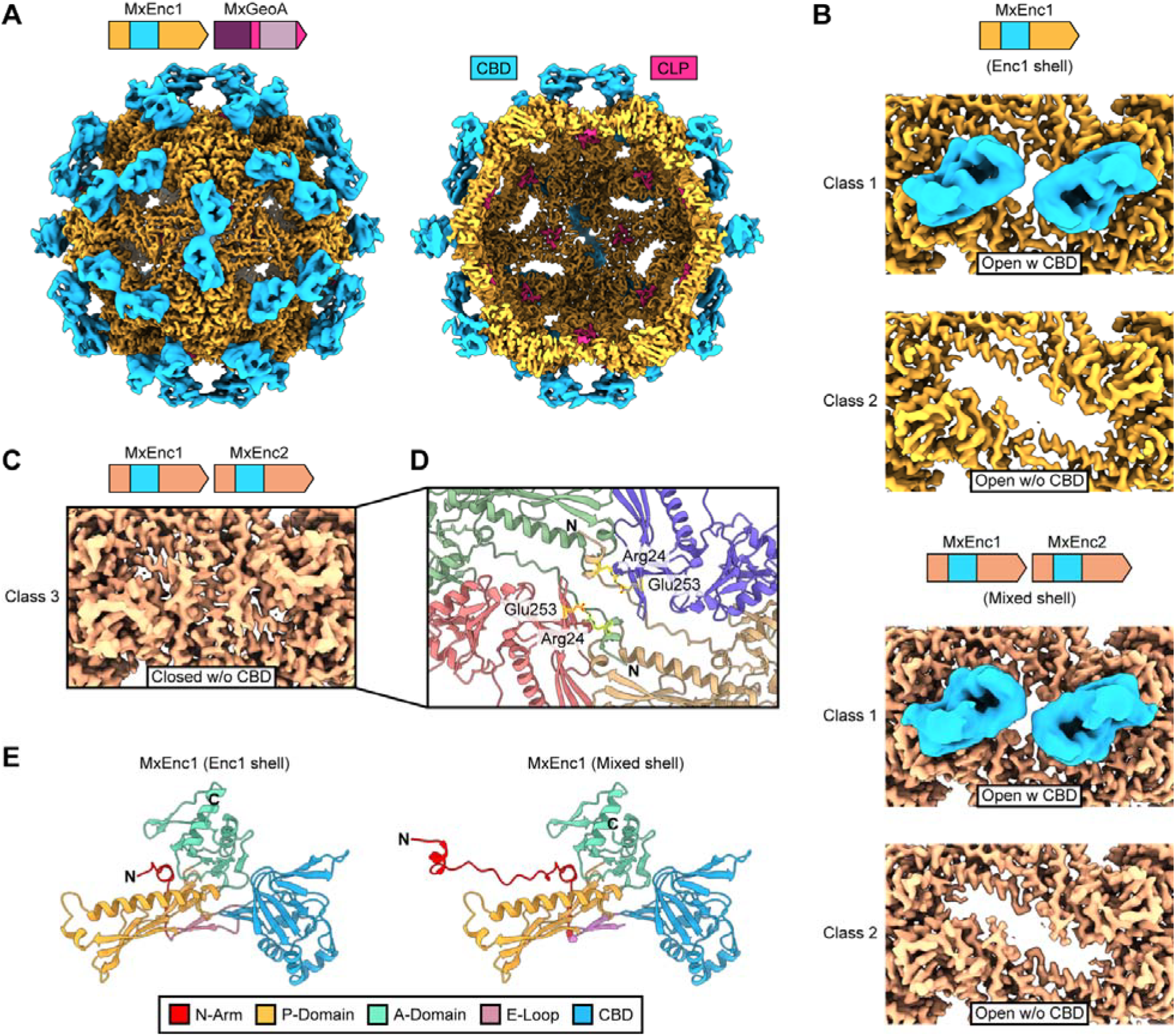
Cryo-EM analysis of GeoA-loaded MxEnc shells. (A) Cryo-EM map of the GeoA-loaded MxEnc1 shell viewed from the exterior (left). Cut-away view of the GeoA-loaded shell (left). Density oft he HK97-like fold is shown in yellow and the external CBDs are highlighted in blue. Cargo CLPs at the internal three-fold axes of symmetry are shown in pink. (B) Zoomed view of the exterior two-fold axis of symmetry in the MxEnc1 shell (top) and MxEnc1Enc2 shell (bottom). Open pores were observed in both shells with CBDs (blue) sitting above the pores at the exterior of the shell. (C) Zoomed view of a third class of two-fold conformation observed in MxEnc1Enc2 shells, where the pore is closed and CBDs are not resolved. (D) Zoomed view of the protomeric N-arms at the closed two-fold pore in MxEnc1Enc2 shells. The N-arms adopt anti-parallel conformations allowing Arg24 and Glu253 to form hydrogen bonding interactions, completely occluding the normally wide open two-fold pore. (E) Structural model of the MxEnc1 protomer in Enc1 only and mixed shells. The characteristic HK97-like domain elements and CBD are highlighted. In MxEnc1Enc2 mixed shells, the N-arm of Enc1 forms a more structured conformation, allowing resolution of the full N-arm structure.

While CBDs could not be directly modeled due to their low resolution, CBD AlphaFold models could be easily docked into the density to create full protomer composite models. Mx CBDs adopt a canonical cAMP-binding domain fold, for example found in *E. coli* catabolite activator protein [50-52]. However, similar to the 2-MIB encapsulin system, no cAMP binding was observed (Figure S10). In native cAMP-binding proteins, ligand binding induces conformational changes that can convey downstream regulatory signals [53, 54]. The same conformational and regulatory logic has been proposed for Family 2B encapsulin CBDs where ligand binding would induce a conformational change resulting in two-fold pore closure, but as yet no CBD ligands have been identified [35].

Masked 3D classifications of the two-fold axis of symmetry yielded different two-fold conformations with respect to resolved/unresolved CBDs and open/closed pore states. Interestingly, the observed two-fold conformations differed between MxEnc1-only shells and mixed shells. The MxEnc1-only shell showed two different types of classes, both with an open two-fold pore with either resolved or unresolved CBDs (Figure 2B). The same classes were also observed in the mixed MxEnc1/MxEnc2 shell, plus a third class with a completely closed two-fold pore without resolvable external CBDs (Figure 2C). This is the first observation of a completely closed two-fold pore in a Family 2B encapsulin shell and confirms for the first time the existence of closed shell states in Family 2B systems, thought to be essential in regulation and control of encapsulated cargo enzymes. The fact that the closed shell state was only observed in mixed shells, containing both MxEnc1 and MxEnc2 protomers, potentially suggests that MxEnc2 is required to achieve a closed conformation around the two-fold pore. The primary difference between open and closed pore states is the absence or presence of structured N-arms that cross the two-fold axis of symmetry, resulting in externally displayed N-termini for two of the protomers directly surrounding the pore. The two respective N-arms adopt anti-parallel conformations which allows the formation of hydrogen bonding interactions between their Arg24 residues and the Glu253 residues, located within the E-loops, of the two neighboring protomers also surrounding the two-fold pore (Figure 2D, 2E). These hydrogen bonding interactions likely stabilize the closed two-fold pore state.

### GeoA Cargo Loading is Mediated by the Cooperative Action of two Conserved Cargo Loading Peptides

Purification of GeoA-loaded encapsulin shells, with the shell composed of either only MxEnc1 or MxEnc1/MxEnc2, resulted in ∼25% of interior three-fold binding sites being occupied with cargo CLPs. In cryo-EM reconstructions, the averaged nature of the CLP density did not permit model building due to limited quality and resolution. Whilst no CLP densities were observed in the absence of cargo as expected, asymmetric C1 refinements in the presence of cargo still yielded, now variable, densities around the three-fold axis binding sites. Due to heterogeneity in CLP binding we were unable to assign amino acid sequences to CLP densities.

The cargo loading domains of characterized Family 2B encapsulin cargo proteins are generally flexible, extended domains, rich in glycine, alanine, and proline [29, 41, 55, 56]. Analysis of the MxGeoA sequence and AlphaFold models showed two stretches of potentially disordered, hydrophobic residues of about thirty amino acids each; one at the intersection between the two GeoA domains (CLP1) and one at the C-terminus (CLP2) (Figure 3A) [27, 57]. Interestingly, the only conserved residues within these stretches were almost identical in CLP1 and CLP2 with the sequence GPN/TGLGTSAAR (Figure 3B, S7, S8). We reasoned that these were the likely peptides mediating shell binding, predominantly through hydrophobic interactions. AlphaFold predictions of binding interactions of CLP1 or CLP2 with the threefold pocket of the MxEnc1 shell showed that both CLP1 and CLP2 could potentially form binding interactions in the requisite binding pocket, as well as with the required interaction geometry, similar to the observed cryo-EM CLP densities (Figure 3C).

**Figure 3:**
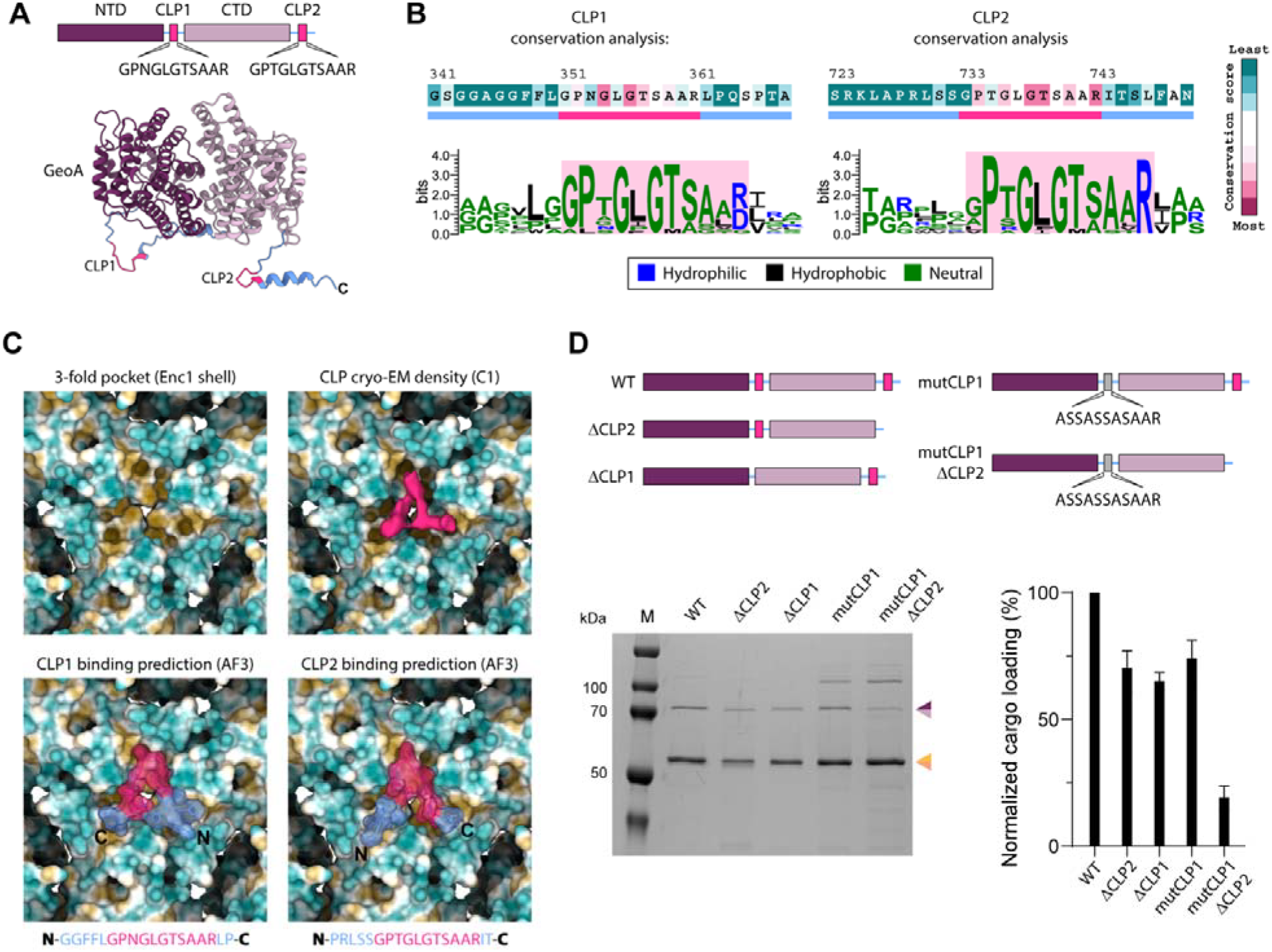
Bioinformatic, structural, and mutational analysis of CLP-mediated cargo loading of MxEnc shells. (A) Schematic overview of GeoA (top) highlighting the N-terminal domain (NTD), C-terminal domain (CTD), internal cargo loading peptide (CLP1), and C-terminal cargo loading peptide (CLP2). AlphaFold predicted structure of MxGeoA (bottom) indicating the structural positions of CLP1 and CLP2. (B)Sequences of MxGeoA CLP1 and CLP2 colored by ConSurf conservation score. A sequence logo of the conserved CLP1 and CLP2 regions is shown, with residues colored by the nature of the amino acid. (C)Zoomed view of the interior three-fold pocket of the MxEnc1 shell (upper left). Zoomed view of extracted cryo-EM CLP density obtained via asymmetric C1 refinement (pink) at the three-fold pocket (upper right). AlphaFold predicted interaction between CLP1 (lower left) or CLP2 (lower right) at the three-fold binding pocket. Both CLPs are predicted to interact at the requisite pocket, and adopt a similar conformation to that observed in cryo-EM reconstructions. (D) Schematic overview (top) of the *geoA* mutants created to test loading of cargo with variable CLP composition. SDS-PAGE analysis of each mutant coexpressed with MxEnc shell proteins and normalized loading percentages based on gel densitometry and using encapsulin expression as an internal control in each case (bottom). Cargo loading is shown as mean values, with error bars representing the standard deviation of three independent experiments.

To further elucidate the role played by each of the putative MxGeoA CLPs, we created a series of deletion and substitution mutants to observe the effects of CLP alteration, both individually and in tandem. Deletion of either the internal CLP1 or the C-terminal CLP2 led to similar but modest, ∼30% reductions in the level of cargo loading (Figure 3D). Mutation of CLP1 to the sequence ASSASSASAAR, also led to a similar reduction in cargo loading. The highest effect on cargo loading was observed when CLP1 was mutated and CLP2 was completely absent, with detected cargo loading levels as low as ∼20% of that seen for WT MxGeoA (Figure 3D). Deletion of both CLP1 and CLP2 led to poor expression and insoluble protein, so cargo loading levels could not be determined for this construct. Taken together this mutational analysis shows that both MxGeoA CLPs, individually, likely play a role in encapsulin binding and cargo loading, and that retention of either peptide is sufficient for cargo loading. In addition to mutational analyses of CLPs in the context of MxGeoA, we also tested minimal peptide fusions to a small test protein, as well as *in vitro* loading of a synthesized minimal CLP peptide (GPTGLGTS). Notably, cryo-EM analysis of the minimal peptide loaded shell also showed heterogeneous CLP densities at the threefold binding pocket, highlighting that flexible, hydrophobicity-mediated binding is likely a feature of the system, and that CLP binding to the interior of the MxEnc shell is mediated by multiple binding modes of CLP1 and CLP2, explaining the difficulty of obtaining high-quality CLP densities via cryo-EM analysis, even after extensive local classifications.

### Encapsulated GeoA Exhibits no Inhibition of Enzymatic Activity Compared to the Free Enzyme and Shows an Improved Geosmin Product Titer

The biosynthesis of geosmin has been studied extensively in both native bacterial hosts, as well as through *in vitro* enzymatic assays [21, 58, 59]. This study is the first report of the biosynthetic enzyme GeoA being enclosed in an encapsulin shell, presenting a number of potential challenges and nuances to effective enzyme activity. The shell represents an obvious diffusion barrier and the effect of an internally tethered GeoA, as opposed to a free, cytosolic, enzyme on enzyme catalysis is unknown [30, 60, 61].

To evaluate the functional viability of the encapsulated GeoA enzyme, the catalytic performance of the complete mixed-shell encapsulin system (EncGeoA) was compared directly to the free, unencapsulated enzyme. *In vitro* EnzChek assays monitoring the initial conversion of FPP to the intermediate (1E,4S,5E,7R)-germacra-1(10),5-dien-11-ol (germacradienol), through detection of released pyrophosphate, revealed identical catalytic behavior between the encapsulated and free enzymes (Figure 4A, S9). The binding affinities and steady-state catalytic parameters also remained consistent for both free GeoA and EncGeoA (*K*_m_ 9.55 and 9.32 µM, respectively and *K*_cat_/*K*_m_ 18535 and 19012 M^-1^ s^-1^). This enzymatic behavior indicates that the porous encapsulin shell does not impose steric hindrance or impede molecular flux, allowing bulk FPP substrate unrestricted entry into the lumenal space.

**Figure 4:**
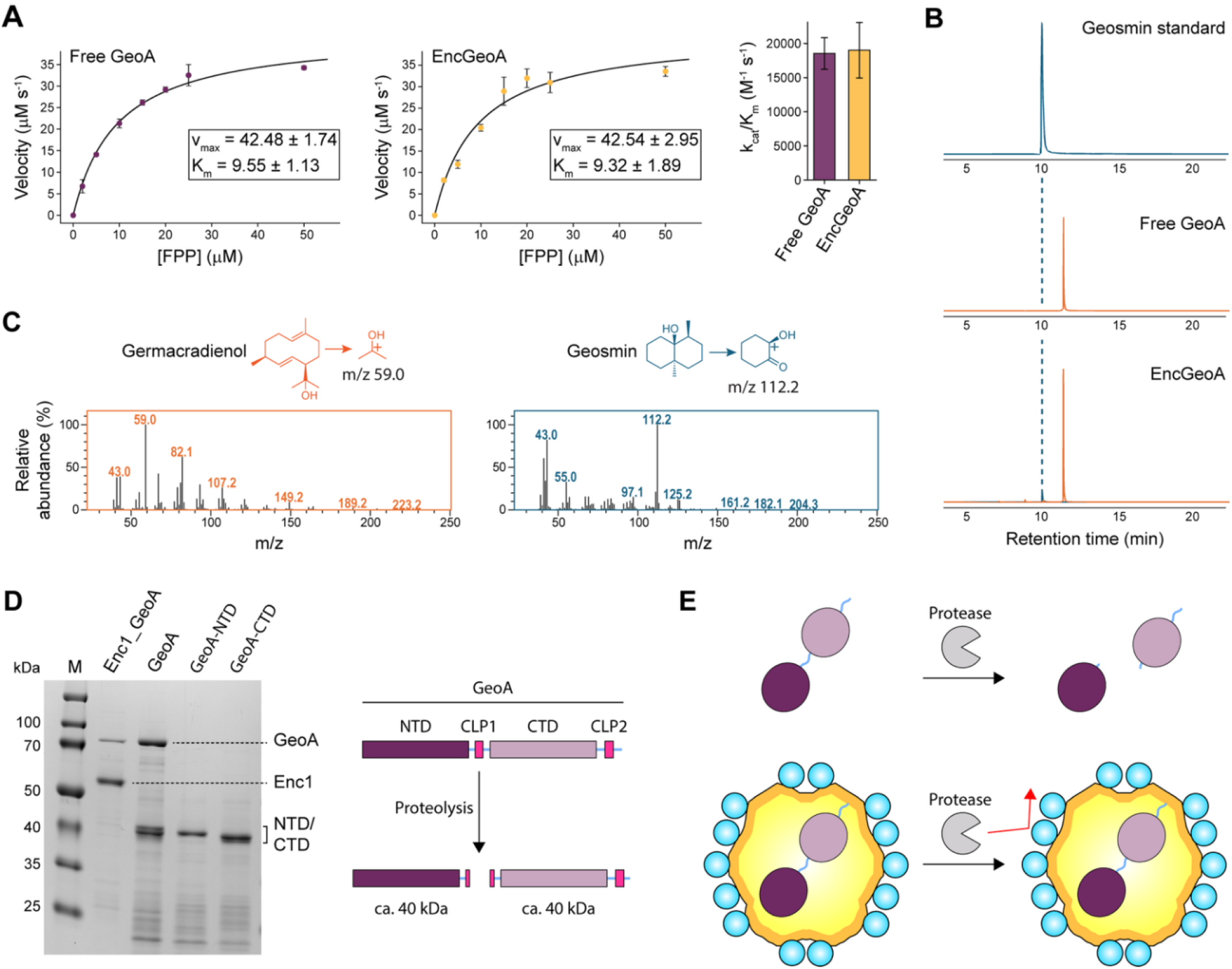
Production and catalytic comparisons of encapsulated and free GeoA enzymes. (A) *In vitro* activity assays of both free and encapsulated GeoA, with the farnesyl pyrophosphate (FPP) substrate. Calculated velocities are plotted as the mean of three independent experiments, with error bars indicating the standard deviation in each case. V_max_ and K_m_ were calculated by applying the Michaelis-Menten equation, which was then used to calculate and plot the catalytic efficiency (K_cat_/K_m_) of the enzyme in each case (right). (B) GC-MS analysis of *in vitro* GeoA activity assays, with both the free and encapsulated enzyme. The intermediate, germacradienol (orange), and product, geosmin (blue), were the main products observed for all assays. Extracted ion chromatograms (EICs) for each of the major products were extracted using diagnostic fragment masses as query m/z (germacradienol – 59.0, geosmin – 112.2). A geosmin standard is shown as a comparison. The y-axis is set to 1 x10^8^ for every chromatogram. (C) Mass spectra collected for each of the major products observed in GC-MS analysis of *in vitro* assays. The expected major fragment masses for both germacradienol and geosmin were observed, confirming the chemical nature of the produced compounds. (D) SDS-PAGE analysis of heterologously purified free and encapsulated GeoA, as well as MxGeoA N- and C-terminal domains. The free enzyme shows break down to smaller mass fragments, resembling the two GeoA domains, at every step of purification, whereas the encapsulated enzyme remains stable throughout prolonged purification and storage protocols. A schematic overview of the likely cleavage point of MxGeoA is also shown (right). (E) Schematic overview of proteolytic propensity of free GeoA (top), compared with the protective phenotype observed when GeoA is encapsulated within a protein shell (bottom).

Subsequent end-point analysis via GC-MS were performed to quantify the overall production of the volatile final sesquiterpene product geosmin. Reactions with both free GeoA and EncGeoA exhibited substantial accumulation of the germacradienol intermediate, as is seemingly characteristic of the enzyme and has been extensively noted in previous *in vitro* studies (Figure 4B, S9) [22, 62]. However, despite identical initial kinetic parameters, EncGeoA demonstrated a notably improved final geosmin product titer compared to the free enzyme (Figure 4B, 4C, S5). We attribute this enhanced overall conversion efficiency to the spatial confinement provided by the encapsulin shell. The shell likely causes a certain level of intermediate trapping and by physically restricting the diffusion of the hydrophobic germacradienol intermediate away from the enzyme active site and into the bulk solvent, the encapsulin maintains a high local concentration of the intermediate within the lumen [28, 63]. This microenvironmental confinement slightly favors the kinetically demanding secondary cyclization and fragmentation step, driving the biosynthesis toward increased geosmin production. Although a large proportion of germacradienol is still formed, similarly to free GeoA, the increase in geosmin formation in encapsulated systems shows a level of intermediate retention and GeoA engagement above the level observed for the free enzyme.

### GeoA Encapsulation Protects the Unstable Free Enzyme from Internal Proteolysis

The *in vitro* utilization of geosmin synthases is historically hampered by severe structural lability and instability; proteins are often purified from insoluble fractions or show increased breakdown [21, 64]. In accordance with previous reports, purification of the full-length free MxGeoA enzyme routinely resulted in significant protein fragmentation, likely driven by internal proteolysis. This observed instability is not unique to the myxobacterial enzyme, but represents a well-documented bottleneck observed across various bacterial GeoA homologs [62]. Analysis of purified MxGeoA by SDS-PAGE, immediately after purification, shows the full-length enzyme, as well as two fragment bands (∼40 kDa) corresponding to the two GeoA enzymatic domains (Figure 4D). The organization of MxGeoA, two catalytic domains connected by a long, flexible linker, leads to long, hydrophobic stretches being solvent exposed [43]. The disordered and exposed ∼30 residues at the intersection of the two MxGeoA domains represent a likely proteolytic cut site. It is unknown whether this proteolysis is an artifact of heterologous expression, but the systemic structural instability observed across multiple GeoA homologs suggests that this feature is likely inherent.

Notably, packaging MxGeoA within the encapsulin shell completely neutralizes this degradation pathway. Purification and extended incubation or storage of EncGeoA results in no degradation of the cargo geosmin synthase. Comparative structural assessments between EncGeoA and MxGeoA demonstrates that the encapsulated enzyme exhibits no detectable proteolytic cleavage, maintaining its full-length integrity where the free enzyme rapidly degrades (Figure 4D). This is likely due to the robust protein shell effectively shielding the vulnerable internal cargo from exogenous proteases preventing unfavorable bulk-solvent interactions that may typically result in instability (Figure 4E) [34]. Beyond enhancing overall product titer, spatial confinement within the encapsulin shell apparently addresses critical structural vulnerabilities inherent to the free enzyme. Consequently, the utility of the encapsulin system extends beyond kinetic advantages and serves an equally vital role as a structural scaffold, conferring long-term stabilization to a notoriously unstable class of terpene synthases.

## Conclusion

In this study, we demonstrate that the two-component Family 2B Mx encapsulin system achieves functional sequestration of the geosmin-forming biosynthetic enzyme GeoA. This is only the second report of encapsulin shell involvement in specialized metabolite biosynthesis and shows that bacterial geosmin formation can be, and in many soil bacteria is, achieved inside a protein shell [41]. Family 2B encapsulin shells have been shown *in silico* to be associated with multiple different types of cargo, including diverse cyclases and polyprenyl transferases, involved in terpene biosynthesis [29]. Additionally, many GeoA-like terpene cyclases were shown to be associated with Family 2B encapsulins, originating from diverse phyla. Interestingly, GeoA-like cyclases were also found to be associated with Family 2A systems, not previously observed to be encoded with terpene biosynthetic cargo, highlighting the diversity of encapsulin/terpene biosynthetic systems.

Biochemical and structural analysis of the complete encapsulin-cargo system showed that both encapsulin protomers are incorporated into the shell, likely in a stochastic manner, as observed for other two-component Family 2B encapsulins [42]. The shells also contained characteristically large two-fold pores, which likely provide the essential flux points for FPP substrate access and geosmin product exit [41, 48]. Interestingly, in comparison to the MxEnc1-only shell, mixed-composition encapsulins displayed an extra two-fold pore state corresponding to a completely closed pore with unresolved external CBDs. This is the first example of a closed two-fold pore in a Family 2B encapsulin system. The now confirmed existence of both open and closed two-fold pore states in Family 2B encapsulins supports the previously hypothesized scenario where CBD ligand binding would result in a conformational change at the two-fold pores, thereby controlling substrate access to the luminal space and ultimately cargo enzyme activity. Closed two-fold pores are formed by specific interactions at the protomeric N-arms, which stabilize the closed state. The N-arms of Family 2B encapsulins show substantial length and sequence variability, which may permit altered orientations of the N-arms and, subsequently, different dynamic two-fold pore states across different systems [35]. As Family 2A encapsulins also generally have near-closed two-fold pores, the coencoding of terpene biosynthetic enzymes with Family 2A shell proteins presents an interesting future line of inquiry regarding substrate and product flux [31, 32, 45].

Sequence analysis of GeoA CLPs yielded two repeating, hydrophobic regions in the internal linker and at the C-terminus. Mutational and structural analysis showed that both of these regions are capable of binding at the internal three-fold axis binding site, highlighting redundant CLP binding capabilities. The limited resolution and observed variability of CLP cryo-EM densities is likely a result of multiple heterogeneous binding modes, as observed in other Family 2B-terpene biosynthetic systems [41]. GeoA homologs encoded alongside Family 2B encapsulins all show extended disordered regions at the C-terminus, as well as at the domain interface, suggesting that this redundant, heterogeneous binding is a conserved feature and likely mediates encapsulation in such systems. Further, almost all GeoA homologs not directly genetically associated with encapsulins also retain these conserved disordered CLP-like regions. 88.3% of the genomes encoding these GeoA enzymes contain at least one, often multiple, other Family 2B genes. This opens up the possibility that GeoA homologs might be encapsulated into Family 2B shells originating from distant genetic loci under certain conditions.

The unique biosynthetic pathway to geosmin has been studied extensively, with the GeoA biosynthetic enzyme analyzed both *in vivo* and *in vitro* [24, 25, 27]. This study shows that the encapsulated GeoA is a functional enzyme, with activity levels on par with the free biosynthetic protein. Encapsulation also increases the titer of the product geosmin, putatively through the process of substrate and intermediate trapping in the lumenal space, causing an increased local concentration of these compounds in proximity to the enzyme active site. This effect has been observed in other encapsulin systems in order to achieve system goals such as maximal storage, or increased activity [65, 66]. Encapsulation also results in protection of the cargo protein from external proteases, as observed by longer cargo protein lifetime compared to the free enzyme. Cargo encapsulation, therefore, provides a stability benefit, in addition to a titer increase, highlighting that the cargo sequestration inside an encapsulin shell can have multiple simultaneous system-specific benefits.

Our results show the extended utility of Family 2B encapsulin shells in bacterial terpene biosynthetic pathways. In addition, we show the first example of a closed two-fold pore in a Family 2B encapsulin system. These intriguing observations open a number of key lines of inquiry, primarily surrounding the role of CBDs in different observed pore states and their role in modulation and regulation of internalized cargo activity. Identifying the native ligand(s) of the MxEnc CBD and the effect binding may have on CBD orientation, and pore state would also be key avenues for future investigation. In sum, our results highlight the multiple functions of a novel two-component encapsulin respect to ligand-mediated cargo enzyme regulation.

