## Supplementary Information for "Bacterial Geosmin Biosynthesis is Compartmentalized Inside a Two-Component Protein Shell"

### **Methods**

**Growth media, strains, plasmids and other reagents**

*Escherichia coli* strains were cultivated using Lennox agar (LA) or lysogeny broth (LB). Culture media was supplemented with antibiotics as required at the following concentrations: ampicillin (100 μg/ml), kanamycin (50 μg/ml). Oligonucleotides and gene fragments were purchased from Integrated DNA Technologies and enzymes were purchased from New England Biolabs unless otherwise stated. All grids and electron microscopy supplies were purchased from Electron Microscopy Services. Reagents for the EnzChek pyrophosphate assay kit were purchased from Thermo Fisher Scientific, and all other solvents and reagents required for various analyses were purchased from MilliporeSigma, unless otherwise stated.

**Computational and phylogenetic analysis**

Analysis of all family 2B encapsulin clusters involved in terpenoid biosynthesis was carried out previously, as detailed [33]. For analysis of all GeoA homologs, a sequence similarity network (SSN) for GeoA was created using the EFI-EST server, initially in summer 2025 [67, 68]. The amino acid sequence of GeoA from *M. xanthus* (Uniprot ID: A0A7Y4MRP8) was used as a BLAST query in EFI-EST with an E-value of −5, a maximum of 5000 sequence hits. This returned 4607 putative GeoA homologs, which reduced to 1627 homologs upon applying a minimum sequence limit of 600 amino acids. The genome neighborhoods of these GeoA homologs were visualized with the EFI-GNT server and classified on the basis of the presence or absence of a second Family 2B encapsulin or a polyprenyl transferase (PPT) encoding gene [67, 68]. All identified clusters were then used to assemble a phylogenetic tree using the NGhylogeny.fr server [69]. This was carried out via MAFFT alignment [70], curation with trimAI and phylogenetic tree assembly with the FastTree algorithm with 1000 bootstraps [71-73]. The tree was visualized and annotated in iTol v6 [74]. Additionally, for SSN analysis of the GeoA proteins within the final dataset, the EFI-EST server was used to assemble an initial SSN, which was then annotated using Cytoscape v3.1.0 to retain only edges with minimally 75% sequence identity [75].

**Molecular cloning**

To produce cargo-loaded or unloaded encapsulin shells in *E. coli*, pET-Duet vectors were utilized, containing either one or both encapsulin-encoding genes (*MxEnc1, MxEnc2*) and with or without the geosmin synthase encoding gene (*geoA*). Each of the genes (*MxEnc1, MxEnc2, GeoA*) were ordered as synthetic gene fragments with twenty nucleotide overlapping sequence, required for Gibson Assembly, after which DNA was used to transform *E. coli* DP10 and plated on ampicillin selective agar plates. Clones were checked for gene insertion by colony PCR, and those containing inserts were whole plasmid sequenced using the Nanopore sequencing service from Eurofins Genomics. Sequenced clones were then used to transform *E. coli* BL21 and used for protein expression and purification. For expression and purification of GeoA in isolation, a pET28a-*geoA* construct was utilized. The coding sequence of *geoA* was PCR amplified using primers containing NdeI and HindIII sites, so that the product fragment contained these sites at the 5’ and 3’ ends, respectively. This insert and the empty pET28a vector were both digested using these same enzymes and the complete plasmid was generated using T4 DNA ligase and used to transform *E. coli* DP10. Clones were confirmed using colony PCR and sequencing and subsequently used for protein expression and purification, as detailed previously.

The mutation of cargo loading peptide (CLP) regions in the *geoA* coding sequence was achieved using specific oligonucleotides in PCR, followed by Gibson Assembly. For any deletion mutants, oligonucleotide primers were used to amplify the *geoA* gene omitting the sequence encoding for the CLP (two overlapping linear fragments for internal deletion, one fragment for C-terminal deletion) and used in Gibson Assembly with the linearized pET-Duet vector. To achieve mutation of the internal CLP, a set of oligonucleotide primers containing the requisite mutations were used to amplify the *geoA* coding sequence, with required overlapping sequence for Gibson Assembly, in two parts. A three-piece Gibson Assembly was performed to clone the mutated *geoA* sequence into the pET-Duet vector, containing the encapsulin-encoding genes.

**Protein expression and purification**

For overproduction of recombinant proteins from *E. coli* BL21, a starter culture of *E. coli* cells harboring the requisite pET-Duet or pET28a plasmid was cultured overnight in 6 ml LB supplemented with either ampicillin or kanamycin while shaking (220 rpm) at 37°C. The complete starter culture was used to inoculate 500 mL of LB media and cultured at 37°C with shaking at 220 rpm, until the cells reached an OD_600_ of 0.6. Protein production was then induced by the addition of 0.2 mM IPTG, followed by shaking at 220 rpm and 18°C for 18 hours. This protocol was utilized for all constructs, apart from pET28a-*geoA* and pET-Duet-*MxEnc1-MxEnc2-geoA*, for which auto-induction media was utilized to optimize yield of protein production. For expression in auto-induction media, a starter culture was again grown overnight in 6 mL LB supplemented with either ampicillin or kanamycin while shaking (220 rpm) at 37°C. The complete starter culture was used to inoculate 500 mL ZY media, supplemented with 1x M and 1x 5052, as described. Cultures were shaken at 220 rpm and at 18°C for 64 hours, after which cells were harvested.

Recombinant protein purification for all constructs was achieved as follows. Bacterial cells were collected by centrifugation and resuspended in lysis buffer (50 mM HEPES, 500 mM NaCl, 10 mM imidazole, protease inhibitor cocktail tablet (Roche), DNAse I, pH 7.5). Cells were lysed by sonication using a Model 120 Sonic Dismembrator (Fisher Scientific Inc.) for three minutes (1s on, 1s off, 84 W power), followed by clarification of the cell lysates by centrifugation (18,000 xg, 20 minutes). The clarified lysates were then passed through a BioRad Econo Pac column loaded with 2 mL Ni-NTA resin (Fisher Scientific Inc.) by gravity flow, before the resin was washed with wash buffer (50 mM HEPES, 200 mM NaCl, 50 mM imidazole, pH 7.5). Protein was eluted from the resin with elution buffer (50 mM HEPES, 200 mM NaCl, 300 mM imidazole, pH 7.5) and analyzed by SDS-PAGE. Elution fractions containing proteins of interest were pooled, concentrated using an Amicon spin filter (Merck, 10 kDA MW cut-off for free enzyme, 100 kDa MW cut-off for encapsulins) and injected on to an ÄKTA Pure system (Cytiva), equipped with either a Superdex 200 increase 10/300 gl (free enzyme) or a Superose 6 Increase 10/300 gl (encapsulin systems) column running a buffer system of 20 mM Tris, 150 mM NaCl, pH 7.5. Resultant fractions from gel filtration showing absorbance at UV280 were collected and analyzed by SDS-PAGE. Fractions containing required proteins were pooled and concentrated using an Amicon spin filter and stored at -80°C, or used immediately.

**Negative stain transmission electron microscopy (TEM)**

Encapsulin protein samples for TEM were diluted to a concentration between 0.1 and 0.15 mg/mL, and applied to freshly glow-discharged (PELCO easiGlow, 60 s, 5 mA) 200 mesh gold formvar/carbon square mesh grids (EMS). Samples were allowed to rest on grids for 60 s to achieve sample adsorption, before liquid was blotted of the grid with filter paper. This was followed by two rounds of staining with 0.75% uranyl formate, achieved by application of a 5 μL drop of the stain and blotting with filter paper. After the application of the second drop of stain, the grid was blotted after one minute to achieve optimal sample staining. TEM micrographs were recorded using either a FEI Morgagni TEM operating at 100 kV or a FEI Tecnai 12 operating at 120 kV, both at the Life Sciences Institute at the University of Michigan.

**Single particle cryo-electron microscopy (cryo-EM)**

Sample preparation: Purified samples of GeoA-loaded His-MxEnc1-only and MxEnc1/His-MxEnc2 mixed shells were concentrated to 0.1 mg/mL and 3 mg/mL in 150 mM NaCl, 25 mM Tris pH 8.0, respectively. 3.5 µL of protein sample were applied to freshly glow discharged grids (MxEnc1-only: Quantifoil R1.2/1.3, 400 mesh, Quantifoil Au holey grid, Graphene coated; MxEnc1/MxEnc2-mixed: Quantifoil R1.2/1.3 Cu 200 mesh) and prepared by plunge freezing in liquid ethane using an FEI Vitrobot Mark IV (100% humidity, 22°C, blot force 0, blot time 4 seconds, wait time 30 s). The grids were immediately clipped and stored in liquid nitrogen until data collection.

Data collection: Cryo-EM movies were collected using a ThermoFisher Scientific Titan Krios G3i cryo-electron microscope operating at 300 kV equipped with a Gatan K3 direct electron detector with a BioQuantum imaging filter. For the Enc1-only sample, 4,596 movies were collected from a single grid using the SerialEM [76] software package at a magnification of 105,000x, pixel size of 0.834 Å, defocus range of -1.0 µm to -1.8 µm, and a total dose of 47.42 e^-^/Å^2^. For the mixed MxEnc1/His-MxEnc2 sample, 861 movies were collected from a single grid using SerialEM at a magnification of 105,00x, pixel size of 0.832 Å, defocus range of -1.0 µm to -1.8 µm, and a total dose of 45.11 e^-^/Å^2^.

Data processing: CryoSPARC 4.7 [77] was used to process the datasets (Supplementary Figs. 4 and 5). For MxEnc1-only, 4,596 movies were imported, motion corrected by patch motion correction, and the CTF fit was estimated using patch CTF estimation. Exposures with CTF fit resolutions worse than 6 Å were discarded from the dataset, resulting in 4,586 remaining movies. 200 particles were selected manually and used to create templates for template-based particle picking. Template picker was then used to select 133,678 particles, which were then extracted using a box size of 480 pixels. Particles subsequently sorted by two rounds of 2D classification, resulting in 121,818 remaining particles. An initial volume was created by ab-initio reconstruction using two classes and I symmetry, resulting in a majority class containing 121,597 particles. These particles were then used for homogeneous refinement against the ab-initio map with I symmetry imposed, per-particle defocus optimization, per-group CTF parameterization, and Ewald sphere correction enabled using a negative curvature sign, resulting in a 2.31 Å map. For MxEnc1/His-MxEnc2, 861 movies were imported, motion corrected by patch motion correction, and the CTF fit was estimated using patch CTF estimation. Exposures with CTF fit resolutions worse than 6 Å were discarded from the dataset, resulting in 861 remaining movies. 200 particles were selected manually and used to create templates for template-based particle picking. Template picker was then used to select 34,983 particles, which were then extracted using a box size of 480 pixels. Particles subsequently sorted by two rounds of 2D classification, resulting in 34,935 remaining particles. An initial volume was created by ab-initio reconstruction using three classes and I symmetry, resulting in a majority class containing 34,935 particles. These particles were then used for homogeneous refinement against the ab-initio map with I symmetry imposed, per-particle defocus optimization, per-group CTF parameterization, and Ewald sphere correction enabled using a negative curvature sign, resulting in a 2.27 Å consensus map. For two-fold pore analysis, particles were symmetry expanded in I resulting in 2,096,200 particles of which 250,000 were used for subsequent 3D classification. Masked 3D classification utilizing a mask encompassing two protomers spanning the two-fold axis of symmetry was carried out with 6 classes, a target resolution of 4.5 Å, O-EM learning rate init: 0.75, and force hard classification enabled. This yielded one class with a closed two-fold pore and unresolved CBDs containing 67,839 particles. Subsequent Homogeneous reconstruction of this class resulted in a closed two-fold pore map with a resolution of 2.71 Å.

Model building: For building the model of the MxEnc1-only shell, a starting model was generated using AlphaFold3 (Q1CYZ6) [78]. The AF3 MxEnc1 model was manually placed into the map using ChimeraX v.1.11.1 [79], followed by improved map fit using the fit-in-map command. The model was then manually refined against the density map using Coot v1.2 [80, 81]. Phenix v2.0.5885 [82, 83] was then used to further refine the model by real-space refinement with three macrocycles, minimization_global enabled, local_grid_search enabled, and adp refinement enabled. NCS operators were then identified from the map using map_symmetry and applied to the model using apply_ncs to generate the icosahedral shell. The NCS-expanded shell was then refined again using real-space refinement with three macrocycles, minimization_global enabled, local_grid_search enabled, adp refinement enabled, and NCS constraints enabled. The BIOMT operators were identified using the find_ncs command and manually placed into the header of the .pdb file of the NCS-refined model (Supplementary Table 1). For generating the closed two-fold model, four MxEnc1 copies were manually placed into the two-fold pore map using ChimeraX v.1.11.1, followed by the fit-in-map command. The model was then manually refined using Coot v1.2, followed by further real-space refinment in Phenix v2.0.5885 with three macrocycles, minimization_global enabled, local_grid_search enabled, and adp refinement enabled.

**GeoA activity assays**

GeoA activity assays were carried out utilizing the EnzChek pyrophosphate assay kit (Thermo Fisher Scientific). Briefly, to assay buffer (50 mM HEPES, 200 mM NaCl, 10 mM MgCl_2_, pH 7.5) was added 2-amino-6-mercapto-7-methyl purine ribonucleoside (MESG), purine nucleoside phosphorylase (PNP) and inorganic pyrophosphatase (IP). This standard mixture was aliquoted at required volumes (total assay volume 100 μL) into wells of 96-well plates, followed by the addition of either free GeoA enzyme or encapsulin-GeoA systems (concentrations ranging from 0 to 4 μM, with encapsulin-GeoA concentrations based on concentration of internal cargo enzyme). Reactions were initiated by the addition of the farnesyl pyrophosphate substrate at varying concentrations from 0 to 100 μM. Absorbance at 360 nm was recorded, corresponding to the conversion of MESG to ribose 1-phosphate and 2- amino-6-mercapto-7-methylpurine by PNP. Readings were collected every 15 s in sweep mode using a BioTek Synergy H1 microplate reader. Slopes were calculated in the linear reaction periods (4 to 10 mins) and used to calculate reaction velocities using a pyrophosphate standard curve. Velocities were plotted and curves were fitted in GraphPad Prism 9.10 or using MatPlotLib.

**Gas chromatography mass spectrometry (GCMS)**

Reactions for subsequent GCMS analysis were prepared in sealed vials to prevent loss of volatile products. In each vial, to assay buffer (50 mM HEPES, 200 mM NaCl, 10 mM MgCl_2_, pH 7.5) was added either free GeoA (4 μM) or EncGeoA (4 μM internal GeoA cargo concentration). Reactions were initiated with the addition of the substrate FPP at concentrations ranging from to 0 to 400 μM to achieve a total assay volume of 100 μL, and incubated for eighteen hours at 30°C. Pentane (200 μL) was added to the vial by syringe to avoid volatile product loss, and mixed by vortexing to transfer pentane-soluble products to the organic layer. The pentane layer was then removed and added to a clean, empty vial. The pentane extraction process was repeated on the aqueous layer to extract any remaining organic compounds. The combined pentane layers were then dried with sodium sulfate and concentrated using a dry nitrogen stream. The pentane extraction was also carried out on control assays containing proteins or substrates only, as well as an assay sample containing 400 μM geosmin standard. 2 μL of each concentrated extract was injected onto a Shimadzu QP-2010 GCMS instrument equipped with a DB-5 column, and run in splitless EI mode. The column flow rate was maintained at 1.0 mL/min with a temperature ramp of 60°C to 280°C at 20°C/min. Analysis of collected data was carried out in GCMSsolution (Shimadzu) and plotted in GraphPad Prism 9.10.

**Differential scanning fluorimetry**

Thermal melts of empty encapsulin, or cargo-loaded encapsulin proteins, was carried out using an Uncle instrument (Unchained Labs). Samples for DSF were either captured using intrinsic tryptophan fluorescence, or contained 10x SYPRO orange dye (from 5000x DMSO stock, Invitrogen), incubated with 1 mg/mL encapsulin protein in storage buffer (20 mM Tris, 150 mM NaCl, pH 7.5). Additional cofactors and binding partners were added into the buffer as required (cAMP, 1mM). Thermal ramps were initiated at 25°C and progressed at 1°C/min, until a final temperature of 95°C was reached. Samples were monitored by obtaining fluorescence emission spectra from 250 to 720 nm, with an excitation wavelength of 473 nm. Data analysis was carried out in Uncle Analysis V5.04, which was used to calculate the area under the curve between 510 and 680 nm and represent thermal transition curves.

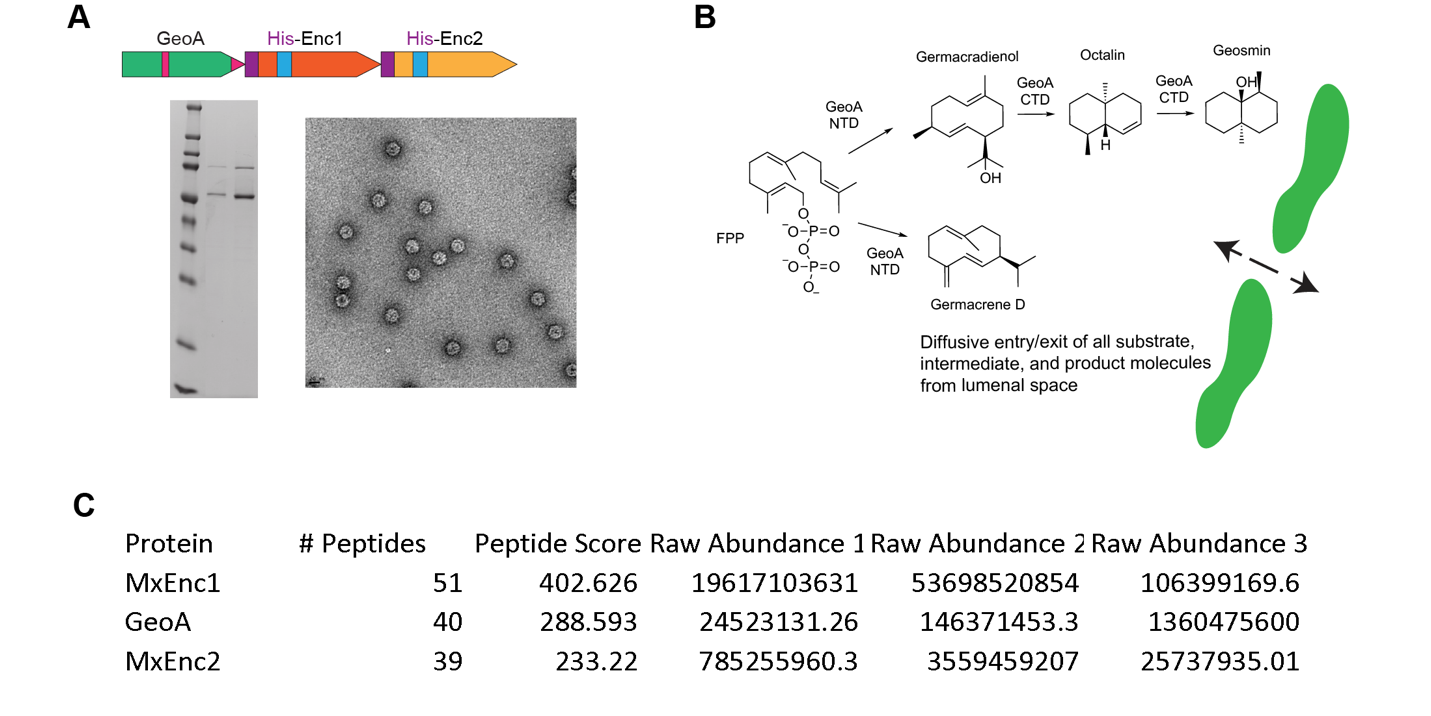

**Supplementary Fig. 1. SDS-PAGE, TEM, and mass spectral analysis of encapsulated GeoA.** (A) Schematic overview of the His-Enc1_His-Enc2_GeoA expressed construct. SDS-PAGE and TEM analysis of the purified protein showed coelution of the shell and cargo proteins and discrete shell formation. (B) Overview of the geosmin biosynthetic pathway and the intermediates formed en route to geosmin. A schematic of an encapsulin pore is shown, illustrating how diffusive substrate and product entry/exit may occur. (C) Results obtained from mass spectral analysis of EncGeoA samples. The overall number of peptides observed across all three samples is shown, as well as averaged peptide scores. The raw mass spectral abundance for each sample is shown for each of the encapsulin proteins and GeoA (samples 1 and 2 – gel slice of encapsulin proteins; sample 3 – gel slice of GeoA protein).

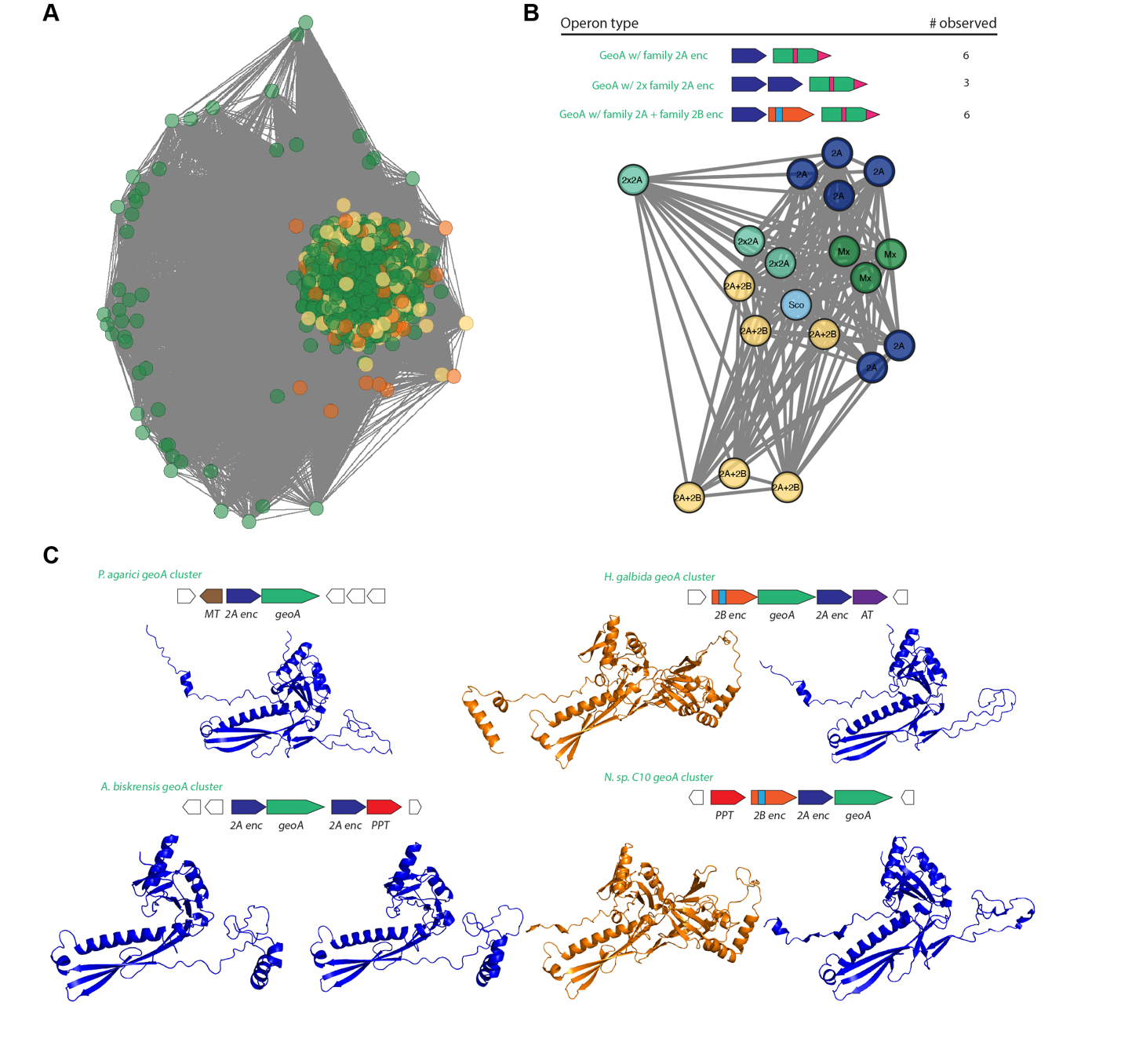

**Supplementary Fig. 2. Sequence similarity networks for GeoA and overview of GeoA systems containing Family 2A encapsulin shells.** (A) Sequence similarity network (SSN) of all analyzed GeoA homologs. Circles highlighted in orange are single encapsulin-containing systems, whereas those highlighted in yellow contain two distinct encapsulin genes. (B) SSN of GeoA systems containing Family 2A encapsulins, as well the GeoA systems from *S. coelicolor* and *M. xanthus* as comparisons. The different Family 2A operon types observed are listed above. (C) Schematic overview of four GeoA-containing clusters found to also encode Family 2A encapsulin genes, as well as AlphaFold analysis of encoded encapsulins. Different system types, either containing one or two encapsulin genes, or either one or both of Family 2A and Family 2B encapsulins, were observed.

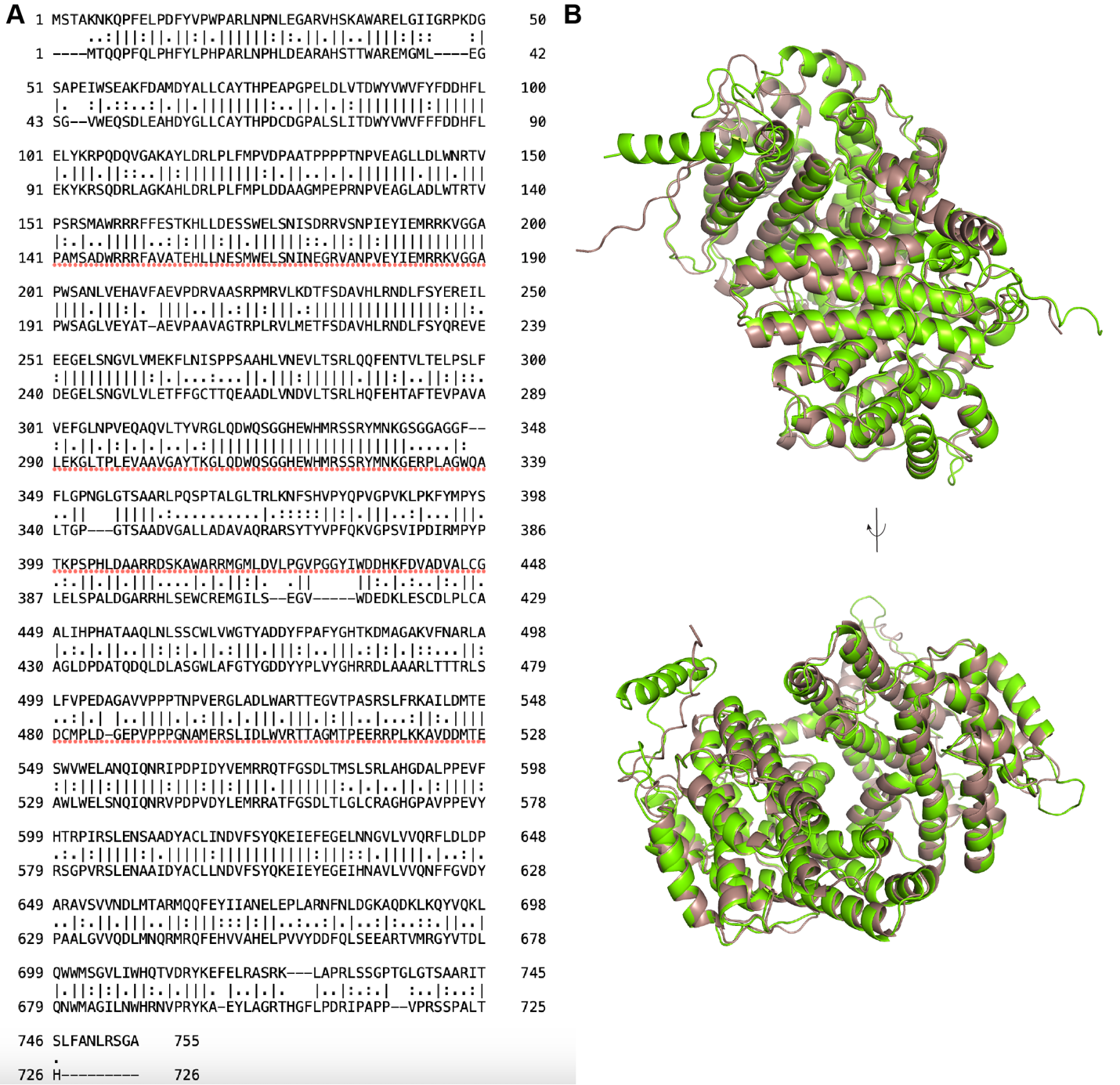

**Supplementary Fig. 3. Pairwise sequence and structural analysis of the GeoA proteins encoded by *M. xanthus* (MxGeoA) and *S. coelicolor* (ScoGeoA).** (A) Pairwise sequence alignment of MxGeoA (top) and ScoGeoA (bottom). The sequences showed 55.5% identity and 70.1% sequence similarity. (B) Alignments of AlphaFold predicted structures for MxGeoA (green) and ScoGeoA (brown). Minor structural differences were only observed at the N- and C- termini of the two proteins.

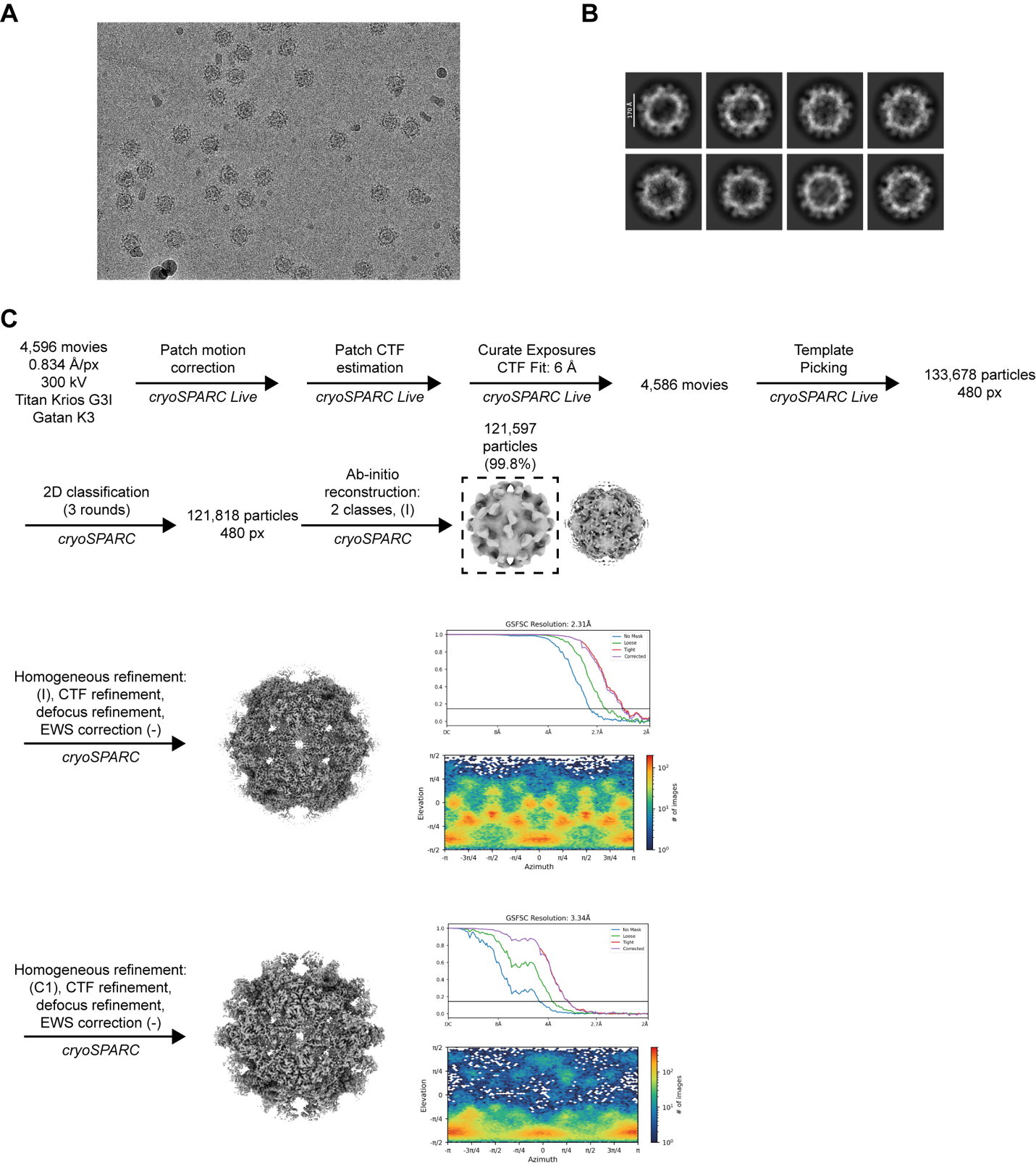

**Supplementary Fig. 4. Cryo-EM workflow for MxEnc1_GeoA.** (A) Representative micrograph for MxEnc1_GeoA. (B) Representative 2D-classes for MxEnc1_GeoA. (C) Overall cryo-EM data collection and analysis workflow carried out for MxEnc1_GeoA.

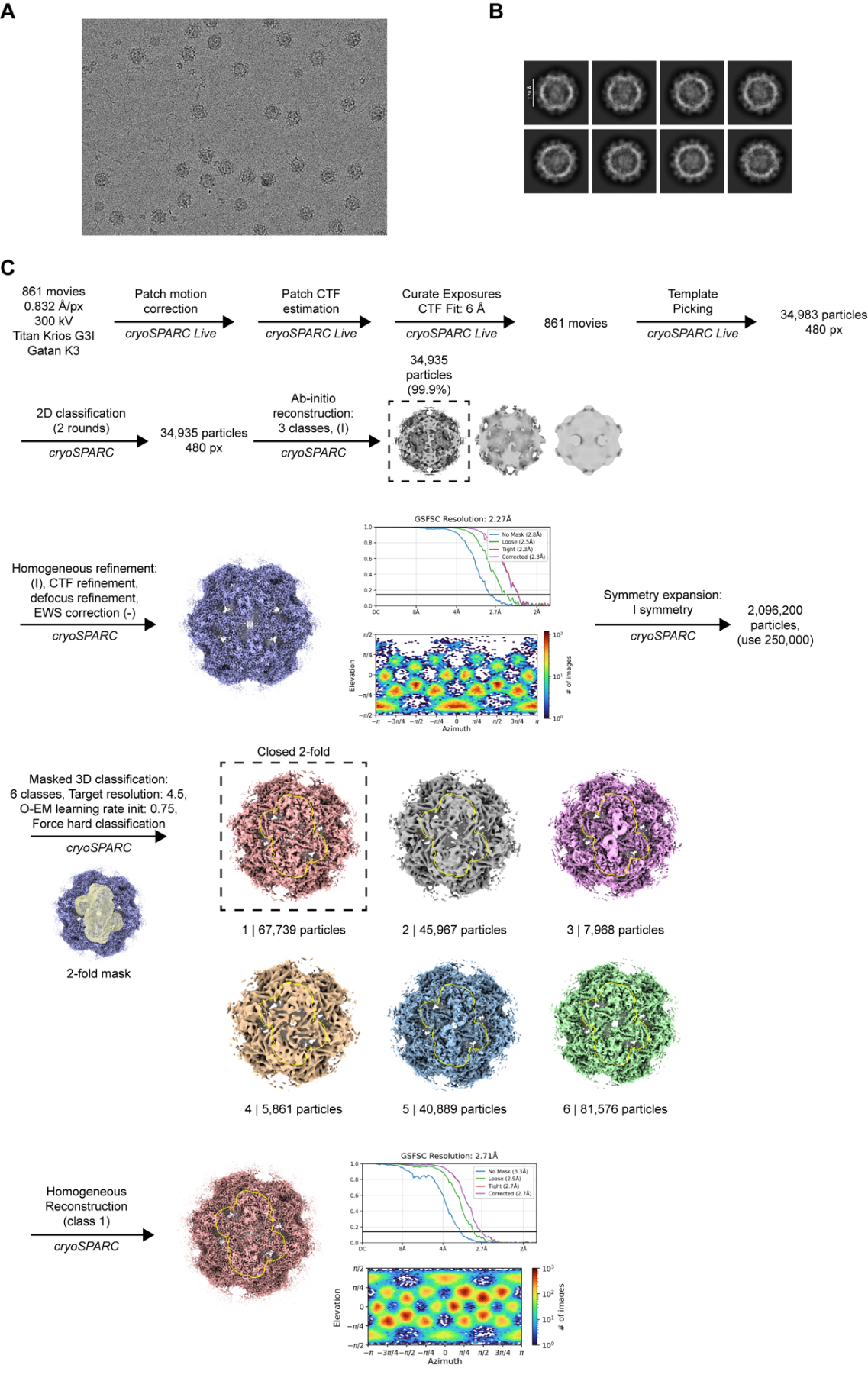

**Supplementary Fig. 5. Cryo-EM workflow for MxEnc1-His-MxEnc2_GeoA.** (A) Representative micrograph for MxEnc1-His-MxEnc2_GeoA. (B) Representative 2D-classes. (C) Overall cryo-EM data collection and analysis workflow carried out for MxEnc1-His-MxEnc2_GeoA.

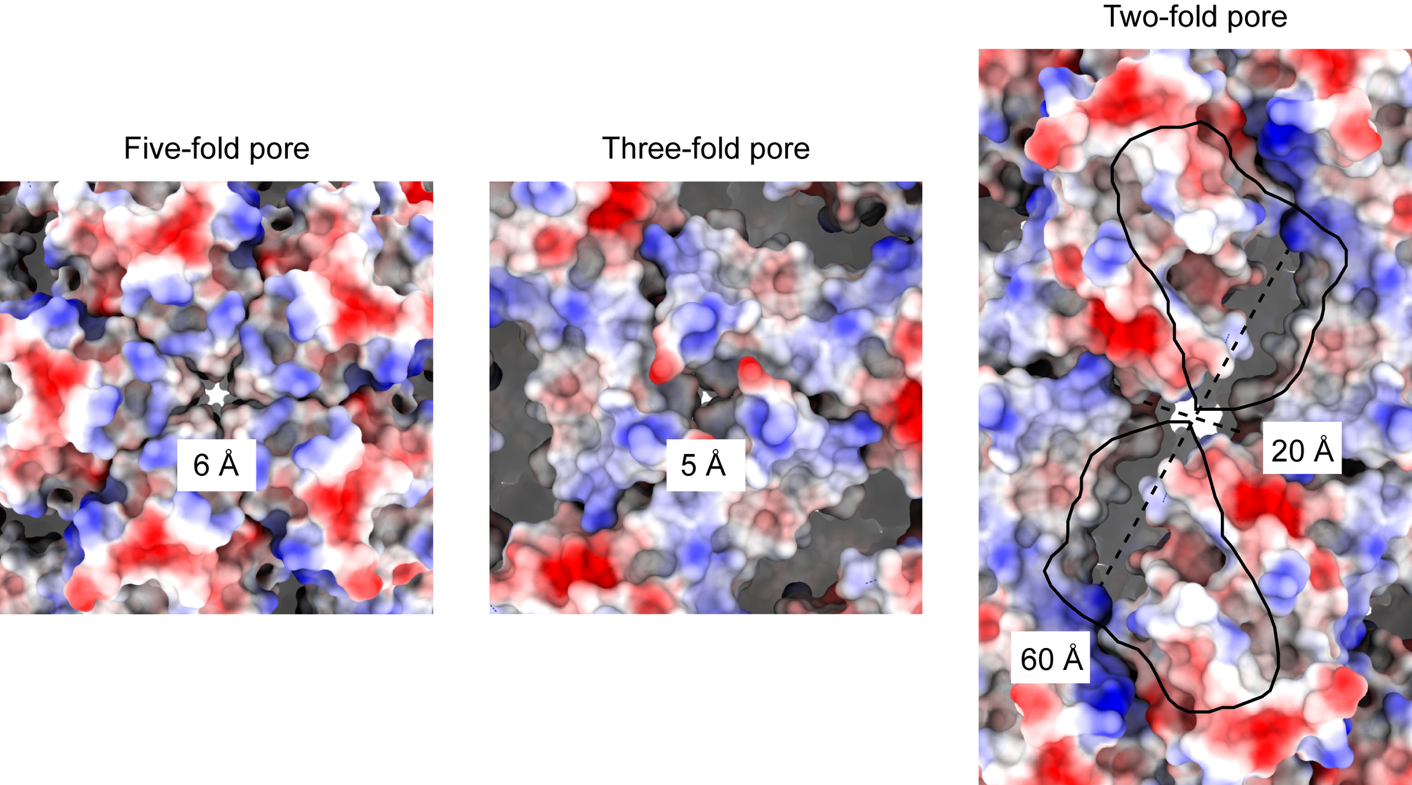

**Supplementary Fig. 6. Structural overview of the three pore types observed in MxEnc encapsulin shells.** (Left) View over the five-fold pore of MxEnc1 shells, colored by charge distribution. The computed size of the pore at its widest is 6 Å. (Center) View over the three-fold pore of MxEnc1 shells, colored by charge distribution. The computed size of the pore at its widest is 5 Å. (Right) View over the two-fold pore of MxEnc1 shells, colored by charge distribution. The external CBDs have been removed for visual clarity, and their outline is indicated above the pore. The computed size of the pore at its widest is ca. 20 Å, with the longitudinal size measured as ca. 60 Å.

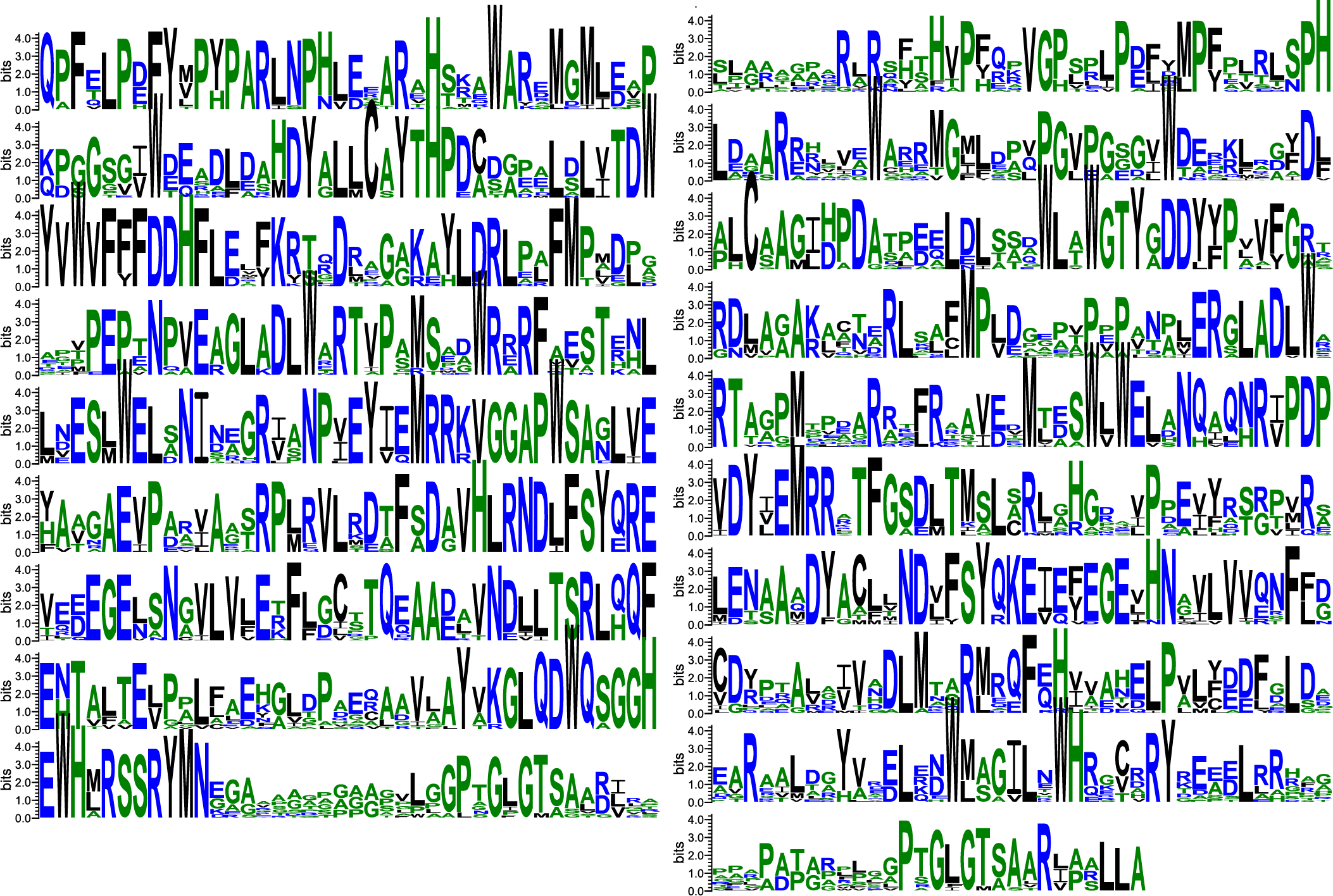

**Supplementary Fig. 7. Computed sequence logo for the complete MxGeoA protein.** The approximate N-terminal domain is shown on the left and the C-terminal domain on the right. Residues in every position are colored by the nature of the amino acid (blue – hydrophilic, black – hydrophobic, green – neutral).

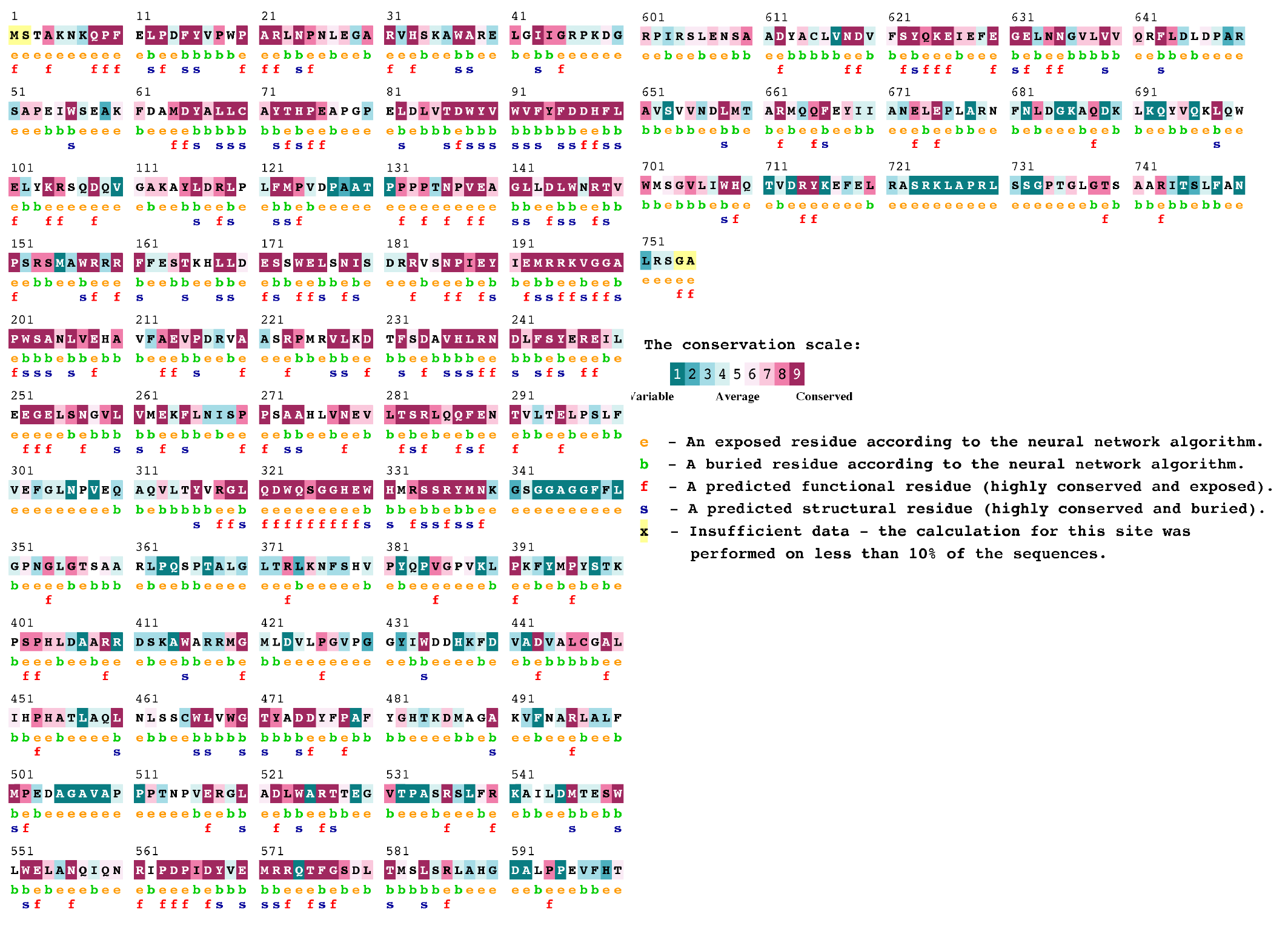

**Supplementary Fig. 8. MxGeoA sequence conservation.** The sequence of GeoA colored by the sequence conservation as computed by ConSurf analysis of 1538 GeoA homologs. Residues are colored and annotated according to the legend on the right.

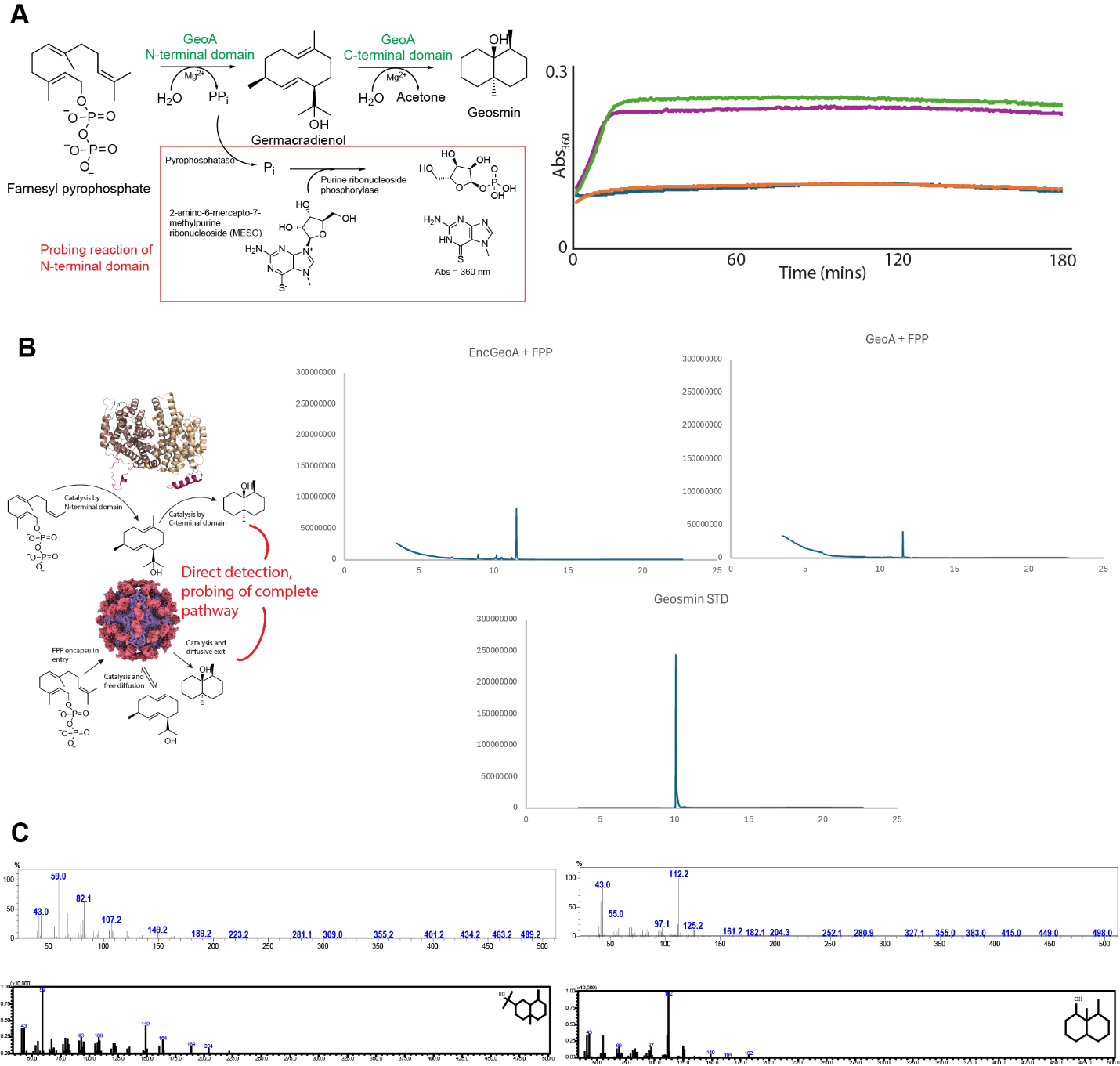

**Supplementary Fig. 9. Catalytic analysis of free and encapsulated GeoA**. (A) Mechanistic overview of the EnzChek assay used to track catalytic activity of both free and encapsulated GeoA enzymes (left). Examples of raw absorbance curves obtained from EnzChek assays with encapsulated GeoA (green (50 μM FPP), 5 μM FPP controls (blue and orange), and free MxGeoA (purple, 50 μM FPP). (B) Mechansitic overview of GCMS assay, used to analyze intermediate and product formation from assays containing substrate FPP and either free or encapsulated GeoA (left). Example total ion chromatograms (TICs) observed in GCMS runs of assays with MxGeoA or EncGeoA with FPP, or a geosmin standard (right). (C) Mass spectra obtained for germacradienol (upper left) and geosmin (upper right) and software-generated comparative spectra for compound assignment confirmation (bottom).

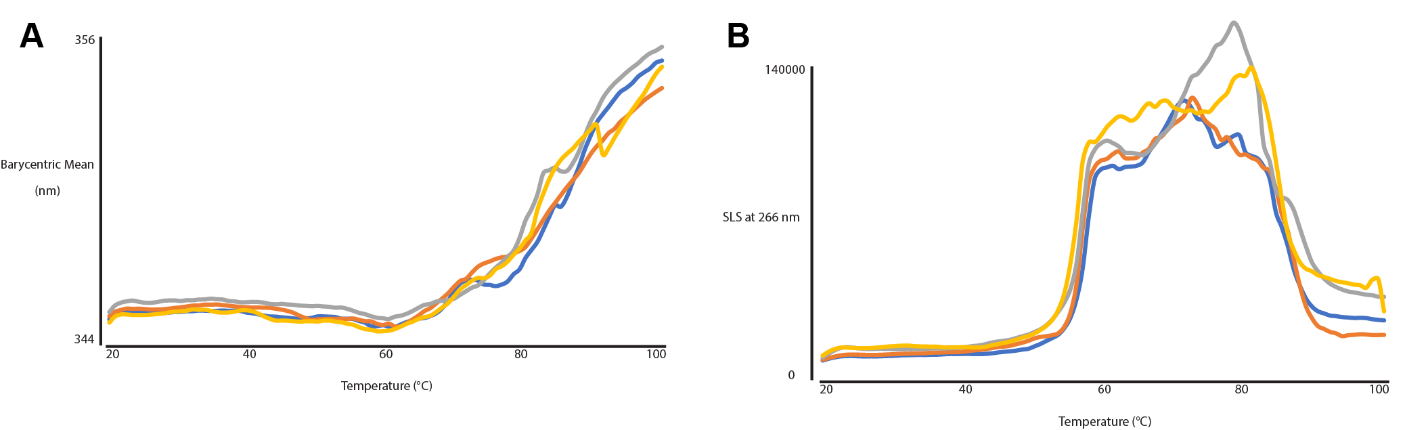

**Supplementary Fig. 10. Differential scanning fluorimetry (DSF) thermal melts of GeoA-loaded MxEnc.** Melt data was collected in the absence (orange, yellow) or presence (blue, grey) of cAMP. Calculated curves from intrinsic fluorescence are shown (left) as well as single light scattering curves at 266 nm (right).

**Supplementary Table 1. Cryo-EM data collection and refinement statistics.**

|  | **Enc1GeoA-I**  **(EMD-77551)**  **(PDB 36GO)** | **Enc1Enc2GeoA-closed 2fold**  **(EMD-77555)**  **(PDB 36GU)** | **Enc1GeoA-C1**  **(EMD-77550)** | **Enc1Enc2GeoA-I**  **(EMD-77552)** |
| --- | --- | --- | --- | --- |
| **Data collection and processing** |  |  |  |  |
| Magnification | 105,000x | 105,000x | 105,000x | 105,000x |
| Voltage (kV) | 300 | 300 | 300 | 300 |
| Electron exposure (e^-^/Å^2^) | 47.42 | 48.23 | 47.42 | 48.23 |
| Defocus range (mm) | -0.8 to -1.8 | -0.8 to -1.8 | -0.8 to -1.8 | -0.8 to -1.8 |
| Pixel size (Å) | 0.834 | 0.832 | 0.834 | 0.832 |
| Symmetry imposed | I | C1 | C1 | I |
| Initial particle images (no.) | 133,678 | 2,098,980 (symmetry expanded) | 133,678 | 34,983 |
| Final particle images (no.) | 121,597 | 275,964 | 121,597 | 34,935 |
| Map resolution (Å)  FSC threshold | 2.31  0.143 | 2.71  0.143 | 3.34  0.143 | 2.27  0.143 |
| **Refinement** |  |  |  |  |
| Initial model used (PDB code) | AlphaFold | AlphaFold | - | - |
| Model resolution (Å)  FSC threshold | 2.5  0.5 | 2.9  0.5 | -  - | -  - |
| Map sharpening *B* factor (Å^2^) | -80.6 | -86.7 | -105.8 | -72.0 |
| Model composition  Non-hydrogen atoms  Protein residues  Ligands | 1,876  237  - | 7,854  1000  - | -  -  - | -  -  - |
| *B* factors (Å^2^)  Protein  Ligands | 44.46  - | 33.18  - | -  - | -  - |
| r.m.s. deviations  Bond lengths (Å)  Bond angles (°) | 0.004  1.055 | 0.008  1.131 | -  - | -  - |
| Validation  MolProbity score  Clashscore  Poor rotamers (%) | 1.95  9.04  2.91 | 1.50  4.88  1.28 | -  -  - | -  -  - |
| Ramachandran plot  Favored (%)  Allowed (%)  Disallowed (%) | 97.4  2.6  0 | 97.05  2.74  0.20 | -  -  - | -  -  - |

| Name  **Supplementary Table 2. Protein sequences of constructs used in this study.** | Protein Sequence |
| --- | --- |
| His-MxGeoA | MHHHHHHGGGSENLYFQGSTAKNKQPFELPDFYVPWPARLNPNLEGARVHSKAWAREL GIIGRPKDGSAPEIWSEAKFDAMDYALLCAYTHPEAPGPELDLVTDWYVWVFYFDDHFLE LYKRPQDQVGAKAYLDRLPLFMPVDPAATPPPPTNPVEAGLLDLWNRTVPSRSMAWRRR FFESTKHLLDESSWELSNISDRRVSNPIEYIEMRRKVGGAPWSANLVEHAVFAEVPDRVAA SRPMRVLKDTFSDAVHLRNDLFSYEREILEEGELSNGVLVMEKFLNISPPSAAHLVNEVLT SRLQQFENTVLTELPSLFVEFGLNPVEQAQVLTYVRGLQDWQSGGHEWHMRSSRYMNKG SGGAGGFFLGPNGLGTSAARLPQSPTALGLTRLKNFSHVPYQPVGPVKLPKFYMPYSTKPS PHLDAARRDSKAWARRMGMLDVLPGVPGGYIWDDHKFDVADVALCGALIHPHATAAQL NLSSCWLVWGTYADDYFPAFYGHTKDMAGAKVFNARLALFVPEDAGAVVPPPTNPVER GLADLWARTTEGVTPASRSLFRKAILDMTESWVWELANQIQNRIPDPIDYVEMRRQTFGS DLTMSLSRLAHGDALPPEVFHTRPIRSLENSAADYACLINDVFSYQKEIEFEGELNNGVLV VQRFLDLDPARAVSVVNDLMTARMQQFEYIIANELEPLARNFNLDGKAQDKLKQYVQKL QWWMSGVLIWHQTVDRYKEFELRASRKLAPRLSSGPTGLGTSAARITSLFANLRSGA* |
| His-MxEnc1 | MHHHHHHSNIIKPGSDAEKSQLSLGTAAARQLATTTKSVPQMQGISSRWLLKLLPWVQVS GGVYRVNRRLSYAVGDGRVTFTTTGAKVQVIPQELCELPLLRSYDDVEVLTALANRFEQK TYKAGDVITEVGKEADCIVLIAHGKVNKIGAGKYGDATVLGVLADGDHYSYEALLESQD YWKFTAKAATASTVLVLQQSDFEAVMAQSPSLHKHVEQFKARSKKKQDTTGQAEIELAA GHTGEPVLPGTYVDYETSPREYELSVAQTVLQIHTRVADLFNEPMNQTEQQLRLTVEALK ERKEHELINNREFGLLHNADLKQRIHTRRGPPTPDDMDELLATVWKEPSFFLAHPRAIAAF GQECSRKGIYPTSVDFNGNMVPAWRGVPIFPCSKIPVSDSRTTSIMLMRAGEKNQGVIGLH PGTIPDEIEAGLNVRFMGINEKAIINYLVTSYFSAAVLVPDALGILESVEIGRGD* |
| His-MxEnc2 | MHHHHHHTKFVKTGDDTPRLSLGTAAARQLATTTKTVPQMQGITPRWLLRMLPWVEVT GGTYRVNRRLSYAVADDRLVFSNIGAKVQVIPQELLKLPLLRGLSGDDEVIAALASQFTQA EYKAGDVIVEAGAPAEHVFLLAHGKARKLRAGPYGAPVTLDVLADGDHFGDQAVVESD DVWTFTVKATTPCTVLSLPQQVFEGLIARSATLRAQVERYRERLKKPQDKLGQAAIPLAA GHSGEPDIQGGYVDYELTPREYELSVAQTVLRVHTRVSDVFSDPMDQTEQQLRLTIEALRE RQEHELINNRDFGLLHNADLKQRVHTRNGPPTPDDLDELLSRRRKSRFFLAHPRTIAAFGR ECTRRGIYPTPIEVQGTPCMAWRGVPLLPCDKIPITEQRTSSILVLRTGQEDQGVIGLRRTGL PDEVEPGLSVRRMDVTDKAITNYLVSTYFSAALLIPDALGVLENVELGG* |
| MxGeoA | MSTAKNKQPFELPDFYVPWPARLNPNLEGARVHSKAWAREL GIIGRPKDGSAPEIWSEAKFDAMDYALLCAYTHPEAPGPELDLVTDWYVWVFYFDDHFLE LYKRPQDQVGAKAYLDRLPLFMPVDPAATPPPPTNPVEAGLLDLWNRTVPSRSMAWRRR FFESTKHLLDESSWELSNISDRRVSNPIEYIEMRRKVGGAPWSANLVEHAVFAEVPDRVAA SRPMRVLKDTFSDAVHLRNDLFSYEREILEEGELSNGVLVMEKFLNISPPSAAHLVNEVLT SRLQQFENTVLTELPSLFVEFGLNPVEQAQVLTYVRGLQDWQSGGHEWHMRSSRYMNKG SGGAGGFFLGPNGLGTSAARLPQSPTALGLTRLKNFSHVPYQPVGPVKLPKFYMPYSTKPS PHLDAARRDSKAWARRMGMLDVLPGVPGGYIWDDHKFDVADVALCGALIHPHATAAQL NLSSCWLVWGTYADDYFPAFYGHTKDMAGAKVFNARLALFVPEDAGAVVPPPTNPVER GLADLWARTTEGVTPASRSLFRKAILDMTESWVWELANQIQNRIPDPIDYVEMRRQTFGS DLTMSLSRLAHGDALPPEVFHTRPIRSLENSAADYACLINDVFSYQKEIEFEGELNNGVLV VQRFLDLDPARAVSVVNDLMTARMQQFEYIIANELEPLARNFNLDGKAQDKLKQYVQKL QWWMSGVLIWHQTVDRYKEFELRASRKLAPRLSSGPTGLGTSAARITSLFANLRSGA* |
| MxEnc1 | MSNIIKPGSDAEKSQLSLGTAAARQLATTTKSVPQMQGISSRWLLKLLPWVQVS GGVYRVNRRLSYAVGDGRVTFTTTGAKVQVIPQELCELPLLRSYDDVEVLTALANRFEQK TYKAGDVITEVGKEADCIVLIAHGKVNKIGAGKYGDATVLGVLADGDHYSYEALLESQD YWKFTAKAATASTVLVLQQSDFEAVMAQSPSLHKHVEQFKARSKKKQDTTGQAEIELAA GHTGEPVLPGTYVDYETSPREYELSVAQTVLQIHTRVADLFNEPMNQTEQQLRLTVEALK ERKEHELINNREFGLLHNADLKQRIHTRRGPPTPDDMDELLATVWKEPSFFLAHPRAIAAF GQECSRKGIYPTSVDFNGNMVPAWRGVPIFPCSKIPVSDSRTTSIMLMRAGEKNQGVIGLH PGTIPDEIEAGLNVRFMGINEKAIINYLVTSYFSAAVLVPDALGILESVEIGRGD* |
| His-MxGeoA NTD | MGSSHHHHHHSSGLVPRGSHMSTAKNKQPFELPDFYVPWPARLNPNLEGARVHSKAWARELGIIGRPKDGSAPEIWSEAKFDAMDYALLCAYTHPEAPGPELDLVTDWYVWVFYFDDHFLELYKRPQDQVGAKAYLDRLPLFMPVDPAATPPPPTNPVEAGLLDLWNRTVPSRSMAWRRRFFESTKHLLDESSWELSNISDRRVSNPIEYIEMRRKVGGAPWSANLVEHAVFAEVPDRVAASRPMRVLKDTFSDAVHLRNDLFSYEREILEEGELSNGVLVMEKFLNISPPSAAHLVNEVLTSRLQQFENTVLTELPSLFVEFGLNPVEQAQVLTYVRGLQDWQSGGHEWHMRSSRYMNKGSGGAGGFFLGPNGLGTS* |
| His-MxGeoA CTD | MGSSHHHHHHSSGLVPRGSHMAARLPQSPTALGLTRLKNFSHVPYQPVGPVKLPKFYMPYSTKPSPHLDAARRDSKAWARRMGMLDVLPGVPGGYIWDDHKFDVADVALCGALIHPHATAAQLNLSSCWLVWGTYADDYFPAFYGHTKDMAGAKVFNARLALFVPEDAGAVVPPPTNPVERGLADLWARTTEGVTPASRSLFRKAILDMTESWVWELANQIQNRIPDPIDYVEMRRQTFGSDLTMSLSRLAHGDALPPEVFHTRPIRSLENSAADYACLINDVFSYQKEIEFEGELNNGVLVVQRFLDLDPARAVSVVNDLMTARMQQFEYIIANELEPLARNFNLDGKAQDKLKQYVQKLQWWMSGVLIWHQTVDRYKEFELRASRKLAPRLSSGPTGLGTSAARITSLFANLRSGA* |
| MxGeoA ΔCLP1 | MSTAKNKQPFELPDFYVPWPARLNPNLEGARVHSKAWAREL GIIGRPKDGSAPEIWSEAKFDAMDYALLCAYTHPEAPGPELDLVTDWYVWVFYFDDHFLE LYKRPQDQVGAKAYLDRLPLFMPVDPAATPPPPTNPVEAGLLDLWNRTVPSRSMAWRRR FFESTKHLLDESSWELSNISDRRVSNPIEYIEMRRKVGGAPWSANLVEHAVFAEVPDRVAA SRPMRVLKDTFSDAVHLRNDLFSYEREILEEGELSNGVLVMEKFLNISPPSAAHLVNEVLT SRLQQFENTVLTELPSLFVEFGLNPVEQAQVLTYVRGLQDWQSGGHEWHMRSSRYMNKG SGGAGGFFLPQSPTALGLTRLKNFSHVPYQPVGPVKLPKFYMPYSTKPS PHLDAARRDSKAWARRMGMLDVLPGVPGGYIWDDHKFDVADVALCGALIHPHATAAQL NLSSCWLVWGTYADDYFPAFYGHTKDMAGAKVFNARLALFVPEDAGAVVPPPTNPVER GLADLWARTTEGVTPASRSLFRKAILDMTESWVWELANQIQNRIPDPIDYVEMRRQTFGS DLTMSLSRLAHGDALPPEVFHTRPIRSLENSAADYACLINDVFSYQKEIEFEGELNNGVLV VQRFLDLDPARAVSVVNDLMTARMQQFEYIIANELEPLARNFNLDGKAQDKLKQYVQKL QWWMSGVLIWHQTVDRYKEFELRASRKLAPRLSSGPTGLGTSAARITSLFANLRSGA* |
| MxGeoA ΔCLP2 | MSTAKNKQPFELPDFYVPWPARLNPNLEGARVHSKAWAREL GIIGRPKDGSAPEIWSEAKFDAMDYALLCAYTHPEAPGPELDLVTDWYVWVFYFDDHFLE LYKRPQDQVGAKAYLDRLPLFMPVDPAATPPPPTNPVEAGLLDLWNRTVPSRSMAWRRR FFESTKHLLDESSWELSNISDRRVSNPIEYIEMRRKVGGAPWSANLVEHAVFAEVPDRVAA SRPMRVLKDTFSDAVHLRNDLFSYEREILEEGELSNGVLVMEKFLNISPPSAAHLVNEVLT SRLQQFENTVLTELPSLFVEFGLNPVEQAQVLTYVRGLQDWQSGGHEWHMRSSRYMNKG SGGAGGFFLGPNGLGTSAARLPQSPTALGLTRLKNFSHVPYQPVGPVKLPKFYMPYSTKPS PHLDAARRDSKAWARRMGMLDVLPGVPGGYIWDDHKFDVADVALCGALIHPHATAAQL NLSSCWLVWGTYADDYFPAFYGHTKDMAGAKVFNARLALFVPEDAGAVVPPPTNPVER GLADLWARTTEGVTPASRSLFRKAILDMTESWVWELANQIQNRIPDPIDYVEMRRQTFGS DLTMSLSRLAHGDALPPEVFHTRPIRSLENSAADYACLINDVFSYQKEIEFEGELNNGVLV VQRFLDLDPARAVSVVNDLMTARMQQFEYIIANELEPLARNFNLDGKAQDKLKQYVQKL QWWMSGVLIWHQTVDRYKEFELRASRKLAPRLSS* |
| MxGeoA mutCLP1 | MSTAKNKQPFELPDFYVPWPARLNPNLEGARVHSKAWAREL GIIGRPKDGSAPEIWSEAKFDAMDYALLCAYTHPEAPGPELDLVTDWYVWVFYFDDHFLE LYKRPQDQVGAKAYLDRLPLFMPVDPAATPPPPTNPVEAGLLDLWNRTVPSRSMAWRRR FFESTKHLLDESSWELSNISDRRVSNPIEYIEMRRKVGGAPWSANLVEHAVFAEVPDRVAA SRPMRVLKDTFSDAVHLRNDLFSYEREILEEGELSNGVLVMEKFLNISPPSAAHLVNEVLT SRLQQFENTVLTELPSLFVEFGLNPVEQAQVLTYVRGLQDWQSGGHEWHMRSSRYMNKG SGGAGGFFLASSASSASAARLPQSPTALGLTRLKNFSHVPYQPVGPVKLPKFYMPYSTKPS PHLDAARRDSKAWARRMGMLDVLPGVPGGYIWDDHKFDVADVALCGALIHPHATAAQL NLSSCWLVWGTYADDYFPAFYGHTKDMAGAKVFNARLALFVPEDAGAVVPPPTNPVER GLADLWARTTEGVTPASRSLFRKAILDMTESWVWELANQIQNRIPDPIDYVEMRRQTFGS DLTMSLSRLAHGDALPPEVFHTRPIRSLENSAADYACLINDVFSYQKEIEFEGELNNGVLV VQRFLDLDPARAVSVVNDLMTARMQQFEYIIANELEPLARNFNLDGKAQDKLKQYVQKL QWWMSGVLIWHQTVDRYKEFELRASRKLAPRLSSGPTGLGTSAARITSLFANLRSGA* |
| MxGeoA mutCLP1ΔCLP2 | MSTAKNKQPFELPDFYVPWPARLNPNLEGARVHSKAWAREL GIIGRPKDGSAPEIWSEAKFDAMDYALLCAYTHPEAPGPELDLVTDWYVWVFYFDDHFLE LYKRPQDQVGAKAYLDRLPLFMPVDPAATPPPPTNPVEAGLLDLWNRTVPSRSMAWRRR FFESTKHLLDESSWELSNISDRRVSNPIEYIEMRRKVGGAPWSANLVEHAVFAEVPDRVAA SRPMRVLKDTFSDAVHLRNDLFSYEREILEEGELSNGVLVMEKFLNISPPSAAHLVNEVLT SRLQQFENTVLTELPSLFVEFGLNPVEQAQVLTYVRGLQDWQSGGHEWHMRSSRYMNKG SGGAGGFFLASSASSASAARLPQSPTALGLTRLKNFSHVPYQPVGPVKLPKFYMPYSTKPS PHLDAARRDSKAWARRMGMLDVLPGVPGGYIWDDHKFDVADVALCGALIHPHATAAQL NLSSCWLVWGTYADDYFPAFYGHTKDMAGAKVFNARLALFVPEDAGAVVPPPTNPVER GLADLWARTTEGVTPASRSLFRKAILDMTESWVWELANQIQNRIPDPIDYVEMRRQTFGS DLTMSLSRLAHGDALPPEVFHTRPIRSLENSAADYACLINDVFSYQKEIEFEGELNNGVLV VQRFLDLDPARAVSVVNDLMTARMQQFEYIIANELEPLARNFNLDGKAQDKLKQYVQKL QWWMSGVLIWHQTVDRYKEFELRASRKLAPRLSS* |

^1^Stops are represented by asterisks

^2^Underlined amino acids represent expression tags
